# *Rēs ipSAE adhūc loquuntur*: What the directionality of predicted aligned error in cofolding models can tell us about scoring diverse molecular complexes

**DOI:** 10.64898/2026.11.02.763449

**Authors:** Roland L. Dunbrack

## Abstract

AlphaFold 3 (AF3) and Boltz-2 estimate confidence in predicted biomolecular complexes using the interface predicted template modeling (*ipTM*) score, which is derived from probability distributions of predicted aligned errors (*PAEs*) between molecular tokens. These errors are directional: the predicted position of tokens in one molecule is evaluated relative to coordinate frames defined by tokens in another. The *PAE* matrix is therefore asymmetric, but AF3 reports the maximum of the two directional pairwise *ipTM* scores, whereas Boltz-2 reports them separately. In both programs, the TM-score scaling parameter, *d*_0_, depends on molecular system size rather than solely on the component being scored. In protein–protein complexes, including disordered regions and domains unrelated to the interface can further lower *ipTM*.

I previously introduced the interaction predicted score from aligned errors (*ipSAE*) to address these limitations for protein–protein complexes. Here, I extend *ipSAE* to interactions involving proteins, nucleic acids, small-molecule ligands, and atomic ions. The framework makes three explicit choices for each molecular pair: which component supplies the alignment frames, which tokens are scored, and how *d*_0_ is defined. For different molecular types, I propose aligning on the more complex molecule and scoring the simpler one. For protein–ligand complexes, this entails aligning on protein residues, scoring all ligand heavy atoms without a *PAE* cutoff, and using a fixed *d*_0_ of 2 Å.

In benchmarks of 102 protein–ligand complexes predicted by AF3 and Boltz-2, this formulation yields a substantially wider range of confidence scores than native *ipTM*. On a held-out set of 82 complexes, thresholds of 0.5 for AF3 and 0.8 for Boltz-2 classified ligand placements with balanced accuracies of 0.938 and 0.862, respectively, using a ligand RMSD threshold of 5 Å. Protein–DNA benchmarks support the same directional scoring scheme, while smaller protein–RNA benchmarks provide a preliminary assessment. These results establish a framework for interpreting predicted structural confidence across different classes of biomolecular interactions and suggest that directional scoring may also be useful for computationally designed protein binders.

The Latin phrase “Rēs ipsae adhūc loquuntur” in the title means “The things themselves are still speaking” referring to the PAE scores and what they tell us about protein/ligand and protein/nucleic acid interactions in cofolding models. The new IPSAE code is freely available from: https://github.com/DunbrackLab/IPSAE2.

## INTRODUCTION

AlphaFold3 (1), Boltz-2 (2), and similar programs can predict the structures of molecular complexes containing proteins, nucleic acids, small-molecule ligands, and metal ions. Molecules are represented as sets of *tokens*, each corresponding to one or more atoms. AF3 and Boltz-2 represent standard amino acids and nucleotides as single tokens and small-molecule ligands as individual heavy-atom tokens. They differ in their treatment of modified amino acids and nucleotides: AF3 generally assigns a token to every heavy atom, whereas Boltz-2 generally retains one token per modified residue. For each token with a valid alignment frame, three reference atoms define a local coordinate system consisting of an origin and three orthogonal axes. For standard amino acids, these frames are defined by the N, Cα, and C backbone atoms, with Cα at the origin and the x-axis bisecting the N–Cα–C angle. The predicted aligned error, *PAE*, estimates the positional error of token *j* that would be observed when the predicted and experimental structures are expressed in local coordinate frames independently defined by token *i*. The *PAE* matrix therefore consists of rows representing the aligned token *i* and columns representing the scored token *j*. Because the reference frames differ when *i* and *j* are exchanged, the *PAE* matrix is inherently asymmetric. For example, aligning on a residue in the rigid core of a protein may produce moderate *PAE* values for residues in a flexible loop. However, aligning on a residue in the flexible loop may produce substantially greater *PAE* values for residues in the core, because uncertainty in the *orientation* of the loop residue’s reference frame is amplified over the distance to the core.

To predict the accuracy of a modeled structure of a single protein, the AlphaFold2 paper introduced the idea of the predicted template modeling score, or *pTM* (3), based on the template-modeling (TM) score formulation of Zhang and Skolnick (4). The TM score was developed to assess the accuracy of predicted models of protein structures compared to experimental structures of the same proteins. After structure superposition, if *d*_*j*_ is a distance between the predicted position of the Cα atoms of residue *j* in the model and residue *j* in the experimental structure of length *L*, the TM score of the model is then defined as:

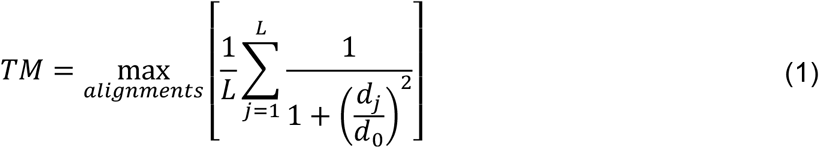

A model of a protein can be superimposed in various ways on an experimental structure, and the maximum is taken over all available alignments (e.g., by running different structure alignment programs or with different parameters or aligning on different segments of the target). The purpose is to identify the largest, well-modeled region of the protein and to minimize the displacements over that region after structural alignment. *d*_0_ is a scaling factor that reduces or eliminates the length dependence of the TM score for alignments of unrelated proteins (4). This is because even completely misfolded models of small proteins cannot have very large *d*_*j*_ values after superposition because the dimensions of the protein are small. Very large proteins can have very large *d*_*j*_ after superposition due to even small changes in intramolecular domain-domain orientations. From a benchmark of unrelated protein pairs of identical lengths (from 40 to 1000 amino acids in length), Zhang and Skolnick fitted *d*_0_ so that unrelated proteins (or very bad models) would have a very similar value of TM (approximately 0.15), regardless of size, which resulted in:

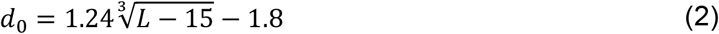

If *L* is 500 residues, *d*_0_ has a value of about 8 (**Figure 1**, left). For longer sequence lengths, the formula for *d*_0_ shifts the TM-score to higher values (**Figure 1**, right). *d*_0_ can also be thought of as the distance *d*_*j*_ that brings the TM score to a value of ½.

**Figure 1.**
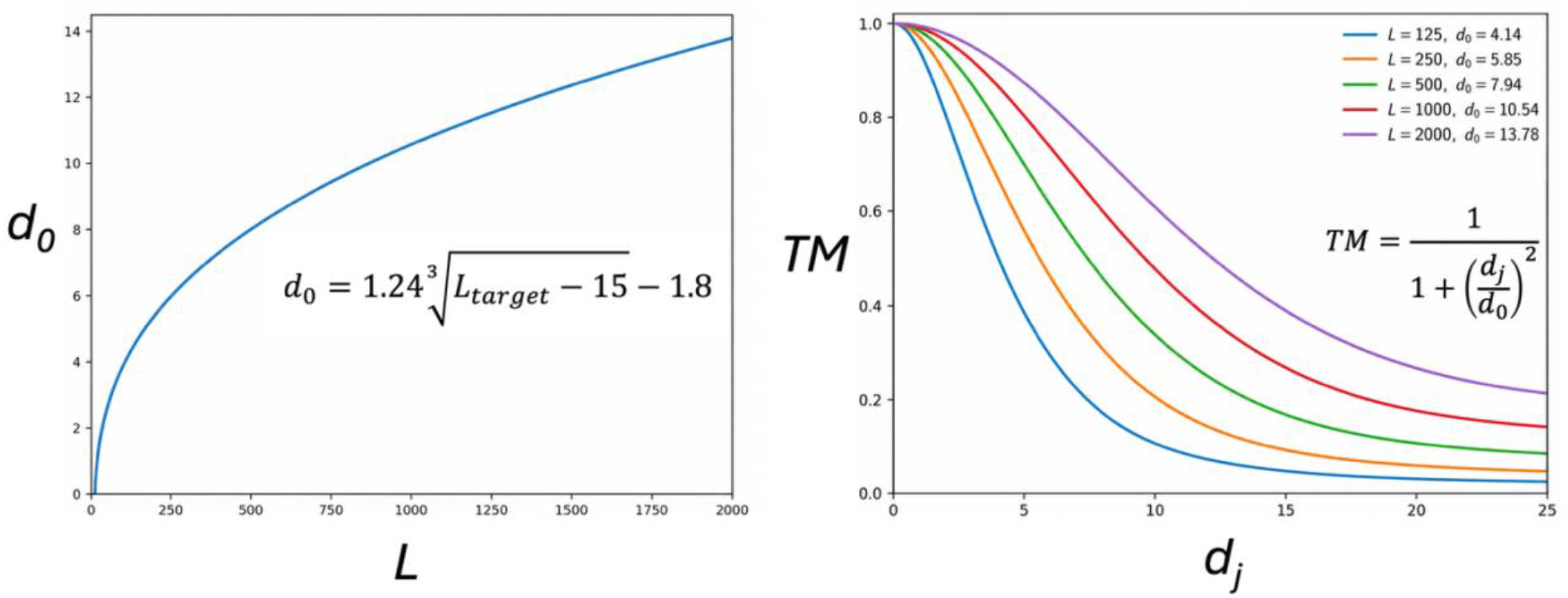
The TM equation: (Left) The *d*_0_ parameter in the TM score equation as a function of target sequence length, *L*. (Right) The function inside the TM score equation as a function of *d*_*j*_ for different values of *L* from 125 amino acids to 2000 amino acids.

In the development of AlphaFold2 (3), a critical insight was that the *PAE*_*ij*_ values produced by the network are analogous to the *d*_*j*_ values in the TM expression (Eq. 1). After structure alignment on residue *i, PAE*_*ij*_ is the predicted uncertainty in the position of residue *j* between the experimental and modeled structures. Thus, a TM score for the structure prediction of a single protein can be directly calculated from the *PAE* matrix by aligning on each residue *i* of the protein, evaluating the function in square brackets in Eq. 1, and taking the maximum over the *L* alignments. In practice, the AlphaFold programs utilize a probability distribution over the aligned error produced by the network, instead of the *PAE* values directly.

With the development of AlphaFold-Multimer (5), the *ipTM* score was introduced for predicting the accuracy of models of protein-protein complexes. For a protein homo- or heterodimer, *ipTM* involves aligning on each residue in one protein and scoring the entirety of the other protein by averaging the *PAE*-probability-based TM score over its full sequence, and vice versa. The output *ipTM* value is the maximum of all such scores across the two proteins. For example, if one protein is 600 amino acids and the other is 900 amino acids, there are 1,500 potential *ipTM* values, and the output score is a maximum of those 1,500 values. For a multi-protein complex, *ipTM* can be calculated pairwise (as in the `chain_pair_ipTM` matrix in AF3 output and `pair_chains_iptm` matrix in Boltz-2 output) or for the whole complex. For the whole complex, each residue in each protein serves as an alignment frame, all the residues in the other chains are scored for that residue, and the maximum over all the alignments is the *ipTM* of the whole complex.

This formulation of *ipTM* creates artifacts that lead to problems in using *ipTM* for protein-protein interaction (PPI) screening. One problem is that the entirety of one protein is scored based on alignment of residues in the other, including the scoring of amino acids in disordered regions and folded domains that are not part of the evolved interface between the two proteins. These regions have large *PAE* values with respect to every aligned residue in the other protein, thus lowering the score significantly. If both proteins have non-binding domains or disorder, then every potential *ipTM* score is lowered, and the final *ipTM* is also lowered. To overcome this, many people break each protein into domains, combinatorially run all domains of one protein against all domains of the other protein and then find the pair with the highest *ipTM*.

The other problem is that AF3, Boltz-2, and other programs calculate the *d*_0_ parameter based on the sum of the lengths of the two proteins (N.B., Boltz-2 actually uses the number of tokens in the entire protein complex, not just in the specific chain pair under consideration). However, the *d*_0_ formula was optimized for the structure prediction of single proteins in which every residue is scored. Calculating *d*_0_ from the sum of the lengths of the two proteins artificially raises the *ipTM* score by scaling the *PAE* values downwards based on the size of the aligned chain and the scored chain jointly. Simply increasing the length of one of the proteins while holding the other fixed increases *ipTM* (as long as the *PAE* matrix remains stable), even if the domain-domain structural interaction remains constant across different-length constructs.

In 2025, I released a preprint presenting a *PAE*-based score called *ipSAE*, which makes up for these deficiencies in the *ipTM* score for protein-protein complexes (6). First, for each residue of the aligned chain, it scores only those residues in the scored protein that pass a *PAE* cutoff with the aligned residue. By using a generous *PAE* cutoff (10 or 15 Å), the score produced is focused on the parts of the scored protein for which there is at least some signal for its placement relative to the other protein. This is usually a single domain or short peptide region that comprises the evolved protein-protein interaction. Thus, disordered regions and folded domains in the scored protein that are completely uncertain in their positions relative to each residue in the aligned protein are not included in the average of the TM function. Second, in *ipSAE*, the *d*_0_ parameter is calculated solely from residues in the scored protein, and only those that are included in the average because they pass the *PAE* cutoff. This formulation more faithfully reflects the origin of the parameterization of the TM score, such that *d*_0_ is based on the size of the scored structure. We and others have shown that *ipSAE* is better at identifying true binding protein pairs from non-binding pairs than *ipTM* (6) and at ranking the structural accuracy of a set of models (6, 7). Other groups have developed similar strategies to transform the *PAE* matrix into a single score for protein-protein pairs, including *LIS* (8) and *actifpTM* (9).

The original *ipSAE* score code (https://github.com/DunbrackLab/IPSAE) outputs the two asymmetric scores for each pair of polymer chains in a predicted structure (e.g., aligning on chain A, scoring chain B; aligning on chain B, scoring chain A) and selects the maximum of those two scores as the *ipSAE* score. The minimum of the two *ipSAE* scores has found utility in filtering computationally designed protein binders (10), although other groups have found the default maximum value better for the same purpose (11). We also included scores for protein/nucleic-acid and nucleic-acid/nucleic-acid pairs. Without much justification, we set the *d*_0_ parameter to 2.0 for complexes involving nucleic acids. Further, the existing code does not provide any scores for ligands or atomic ions.

In this paper, we present a rationale and framework for extending *ipSAE* to interactions of proteins with ligands, atomic ions, and nucleic acids. We continue to provide the two asymmetric scores for each molecule pair. For same-type molecule pairs (protein/protein, nucleic-acid/nucleic-acid, or ligand-ligand), the *ipSAE* score is the maximum of the two asymmetric scores. For different-type molecule pairs, however, we argue that one direction should be preferred over the other. Generally, aligning on the more complex molecule (proteins over nucleic acids, ligands, and ions; nucleic acids over ligands and ions) is the more rational choice and we show that it results in more discriminative scores for ranking structural accuracy. So instead of identifying *ipSAE* as the maximum of the two asymmetric scores for different-type molecule pairs, we define *ipSAE* as the appropriate directional score for each molecular pair type (e.g., aligning on protein residues and scoring ligands and ions).

Two other factors need to be defined for the scored molecule: which tokens to include in the TM score average and what value of *d*_0_ to use. We argued previously that for proteins, it makes sense to score only residues that have decent *PAE* values for each aligned residue in the other chain and to define *d*_0_ based on the number of scored residues in the average. However, when scoring ligands, it makes far less sense to use a *PAE* cutoff since we are interested in scoring the whole ligand in a protein binding site. In contrast to proteins where some parts of a protein may have no well-scored interaction with another protein (disorder or domains not involved in the evolved interaction of the two proteins), there is no part of the ligand that is “irrelevant.”

We also argue that the value of *d*_0_ for scoring a ligand should not be based on the formula of Zhang and Skolnick which was predicated on the alignment of unrelated same-length proteins of different sizes. For protein-ligand *ipSAE*, we hypothesize that *d*_0_ should be a small constant. From a benchmark of 102 protein-ligand targets from the *FoldBench* data set (divided into a training/validation set of 20 targets and a testing set of 82 targets), we determine that a value of *d*_0_=2.0 Å results in a wide dynamic range and effective separations of good and bad models with scoring cutoffs of *ipSAE*=0.5 for AlphaFold3 and *ipSAE*=0.8 for Boltz-2. Nucleic acids are larger and more complicated than ligands, and the choices for scored tokens and *d*_0_ are less clear. Through a small benchmark of protein/nucleic acid complexes, we settle on scoring tokens in the nucleic acid that pass a *PAE* cutoff, since very large nucleic acids may have regions that are not well predicted relative to any residue in a binding protein. We show that defining *ipSAE* as the protein → nucleic acid direction for aligning and scoring tokens respectively and setting *d*_0_ = 2 Å results in a wide dynamic range and effective scoring of accurate vs inaccurate structure predictions.

In the next section, we review the derivation of the *ipTM* score and the motivation and rationale for defining *ipSAE* scores for various molecular interaction categories. After that, we provide some benchmarking data that validate these choices.

## METHODS

### Definition of molecular tokens and the PAE matrix

AlphaFold3 and Boltz-2 divide molecular structures into tokens that are used to model relationships among the components of the entire structure (1). Each standard amino acid or nucleotide is a single token. In AlphaFold3, each modified or non-standard amino acid contains one token per heavy atom, while in Boltz-2 each one is a single token. Most small-molecule ligands contain one token per heavy atom in both programs. Each alignable token *i* is associated with a set of three frame atoms (*a*_*i*_, *b*_*i*_, *c*_*i*_) that define an oriented reference frame with atom *b*_*i*_ as the origin. For standard amino acids in both AF3 and Boltz-2, the three atoms are (N, Cα, C), with Cα as the origin. For standard nucleotides in both programs, the three atoms are (C1′, C3′, C4′), with C3′ as the origin of the reference frame. For atom-level tokens in small-molecule ligands and non-standard residues, a reference frame is constructed using the token atom as the origin and the two nearest heavy atoms in the same molecular component. Tokens without three suitable non-collinear atoms, such as monatomic ions or atoms in small linear molecules (e.g., CO or O_2_), cannot define valid alignment frames.

AF2, AF3, and Boltz-2 use the concept of *aligned error* to generate predicted accuracy metrics for output models (**Figure 2**). For each alignable token *i*, the coordinates of the predicted and experimental structures are expressed separately in the oriented reference frames, centered at the origin, defined by token *i* in each structure (with atom *b*_*i*_ at the origin). For every target token *j*, the distance between the position of a representative atom in the experimental structure and the position of the same atom in the model structure, expressed in their respective reference frames, is calculated. This is the *aligned error*. For amino acids, the Cα atom is used as the representative atom whose position is evaluated. For nucleotides, although C3′ is specified as the frame origin in the AlphaFold3 Supplementary Information, C1′ is the representative atom used to evaluate the aligned error. In contrast, the Boltz-2 source code indicates that C3′ serves as both the frame origin and the atom whose positional error is evaluated. For ligands, ions, and most non-standard amino acids and nucleotides, each heavy atom is a token and serves as its own scored representative atom, even when it cannot define an alignment frame.

**Figure 2.**
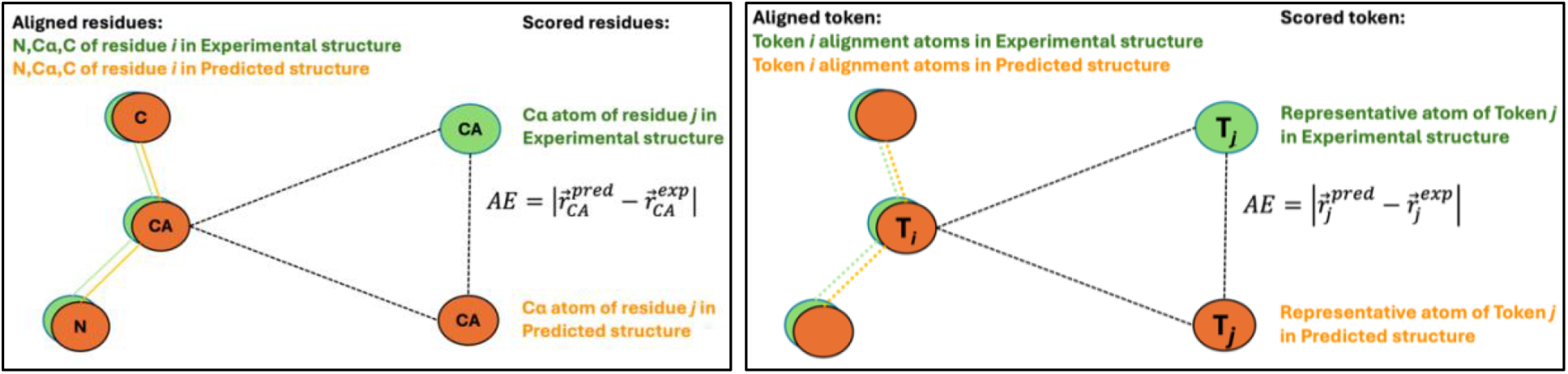
Definition of aligned error (AE) for amino acid pairs (left) and for generalized tokens (right). For generalized tokens, the atom at the origin of the alignment frame (marked *T*_*i*_) is not necessarily the same as the scored or representative atom (marked *T*_*j*_). For tokens in ligands and nucleic acids, the alignment atoms are not necessarily covalently bonded to the token origin atom (dashed orange and green lines).

During training, the experimental structure is known, and the network is trained to predict a probability distribution over the aligned error distance when the experimental structure is not known (i.e., during inference). The probability distribution is defined over the *AE*_*ij*_ distance in 64 bins of width 0.5 Å (0 Å - 0.5 Å, 0.5 Å - 1.0 Å, …, 31.5 Å - 32 Å), where the last bin also includes distances larger than 32 Å. The predicted aligned error, *PAE*_*ij*_, for each pair of tokens is calculated from the predicted probability distribution over the aligned error with the equation (Eq. 11 in AF3 paper supplemental):

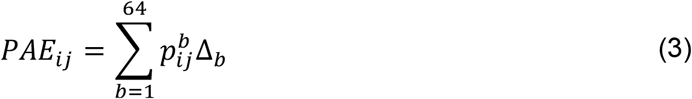

where Δ_*b*_ = (*b* − 0.5)/2 is the center of each bin (0.25 Å, 0.75 Å, …, 31.75 Å), 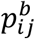 is the probability of bin *b*, and 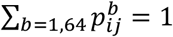.

### Definition of pTM *and* ipTM scores in AlphaFold3 and similar programs

Since the *PAE* values are exactly analogous to the residue displacements after structure superposition in the TM expression (Eq. 1), the probability distribution of the *PAE*_*ij*_ output by the networks can be substituted into Eq. 1 to provide an equation for the *pTM* score (predicted template modeling score) for a single chain (or a whole protein complex):

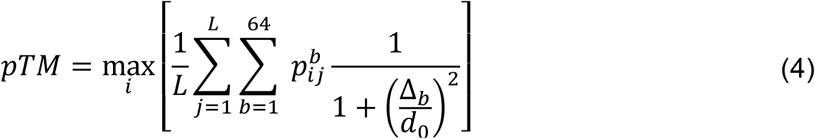

The expression under the sum over distance bins *b* is the expectation value from the probability distribution of the aligned error used in Eq. 3 (AF3 paper supplemental, Eq 17), where Δ_b_ are the bin centers (0.25 Å, 0.75 Å, …, 31.75 Å). The outer average is over all tokens, *j=*1,*L*, in the chain (or structure) and the maximum is taken over all alignable tokens *i*=1,*L* in the structure (i.e. excluding atomic ions or linear molecules). The scored tokens can be from protein residues, nucleotides in DNA or RNA, ligand heavy atoms, ions, or heavy atoms in modified amino acids or nucleotides. The parameter *d*_0_ is taken from Eq. 2 where *L* is the number of tokens (not residues) in the structure. **The role of token *i* in this equation is to create a set of alignments used to calculate the TM score, one for each token in the chain (or complex)**. The value of *pTM* is then calculated from the highest scoring of these alignments, just as in Equation 1 for the original TM score

An approximation to the *pTM* score can be obtained by simply using the *PAE*_*ij*_ themselves:

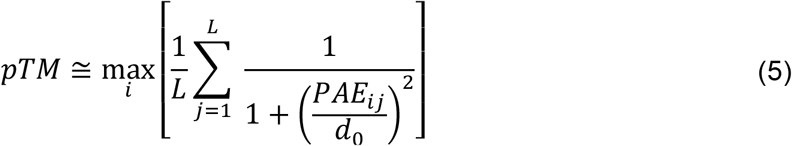

For simplicity, we define the pairwise **pTM** matrix from the aligned error probability distribution as:

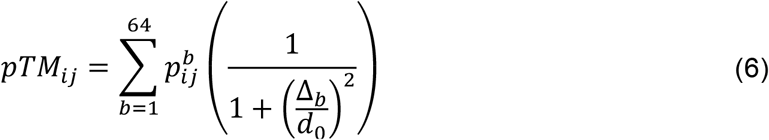

or alternatively (as an approximation) from the *PAE* value.

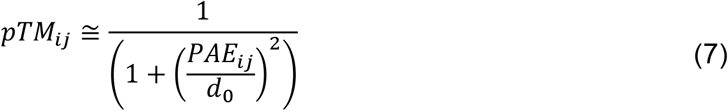

As in the original *ipSAE* implementation, we use the expected *PAE* values rather than the full aligned-error probability distributions, allowing *ipSAE* to be calculated directly from standard prediction outputs. Because the TM function is nonlinear, this is an approximation to the probability-weighted formulation used in native *ipTM*. The approximation in Equation 7 can be substituted for the probability-weighted expression in Equation 6 in subsequent score calculations.

The token-specific mean value of *pTM*_*ij*_ , based on the alignment of token *i* is given by:

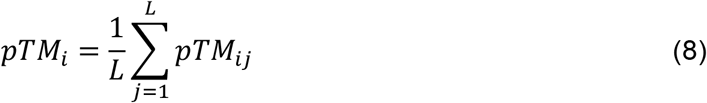

so that *pTM* = max[*pTM*_*i*_].

From these equations, we can generalize the expression for *pTM* by specifying the token sets for the alignments (set *S*_1_, tokens *i*) and those for the token displacements between modeled structure and experimental structure, if it were known (*S*_2_: residues *j*):

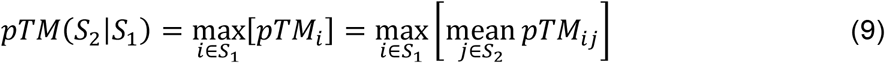

*S*_1_ and *S*_2_ can be different parts of the same chain, such as two domains within a single protein or a core region of a protein domain and a large flexible loop (12). For a complex of two molecules (two proteins, protein and ligand, protein and nucleotide chain, etc.), A and B, we can use tokens in one molecule (e.g., *S*_1_=molecule A) as alignment frames, and calculate the TM score over tokens in the other molecule (*S*_2_=molecule B), which would then contain a rotation-translation component as well as the accuracy of the structural model of molecule B when it is bound to molecule A. So,

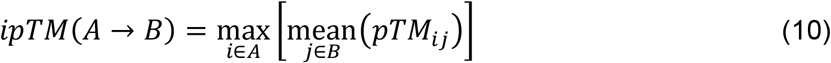

When AlphaFold2 and AlphaFold3 provide a value of *ipTM* for a pair of molecules, they provide a single value which is the maximum of the two asymmetric values (or equivalently the maximum over all alignable tokens in both chains of the interchain *pTM*_*i*_):

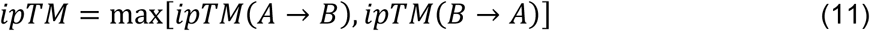

AlphaFold2 and AlphaFold3 use a value for *d*_0_ based on the sum of the total number of tokens in the two molecules. Boltz-2 calculates *d*_0_ from the number of tokens in the entire complex, rather than the pair for each pairwise *ipTM*. This differs from the AlphaFold choice when there are more than two molecules.

AlphaFold3 provides an *ipTM* for each molecule where the maximum is taken over all tokens *i* in that molecule and the mean is over all tokens in all other chains.

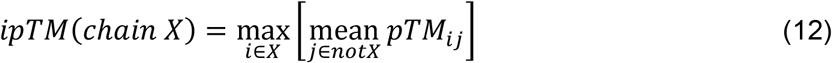

In Eq. 12, *d*_0_ is calculated from the total number of tokens in the model. In AlphaFold2 and AlphaFold3, the overall *ipTM* of any molecular complex is calculated from the maximum over all alignable tokens of the mean *ipTM*_*ij*_ , where the mean is taken over all tokens in all other chains that do not contain token *i*. The value of *d*_0_ is also calculated from the total number of tokens in the model:

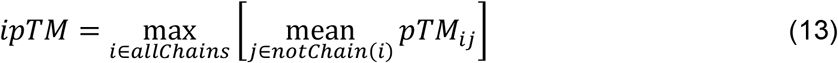

AlphaFold-Multimer does not provide pairwise *ipTM* scores for a multiprotein complex, while AlphaFold3 provides the symmetric `chain_pair_iptm` matrix in the summary confidences file, so that Element [0,1] = Element [1,0] of the AF3 *ipTM* matrix. Boltz-2, by contrast, provides an asymmetric `pair_chains_iptm` matrix. Counterintuitively, Element [0,1] of the Boltz-2 *ipTM* matrix is the value for aligning on tokens in molecule 1 and scoring the tokens in molecule 0.

### Definition of ipSAE for all molecule pairs

We define the generic **asymmetric** *ipSAE* score (*i*nteraction *p*redicted TM *S*core based on *A*ligned *E*rrors) for any two molecules (A and B) as follows:

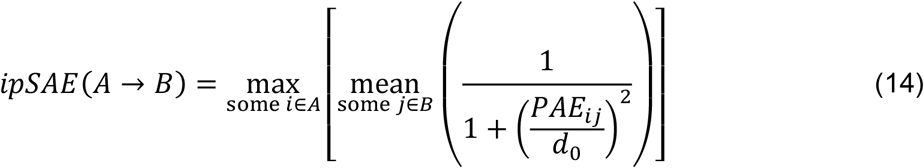

The mean is taken over some or all tokens in molecule B and the max is taken over some or all alignable tokens in molecule A. For every molecule pair, A and B, we can calculate two asymmetric scores: *ipSAE*(*A* → *B*) and *ipSAE*(*B* → *A*). For both A and B, we specify that only some of the tokens in each molecule need to be considered. For instance, we could choose two different domains of the same protein chain, and A and B would represent the same chain.

As we defined *ipSAE* for protein-protein interactions previously, when B is a protein, we use a *PAE* cutoff to avoid the inclusion of regions of chain B that are disordered or found in domains that are not part of the interaction. We can do this whether A is a protein, nucleic acid chain, or ligand. So, when B is a protein, called P:

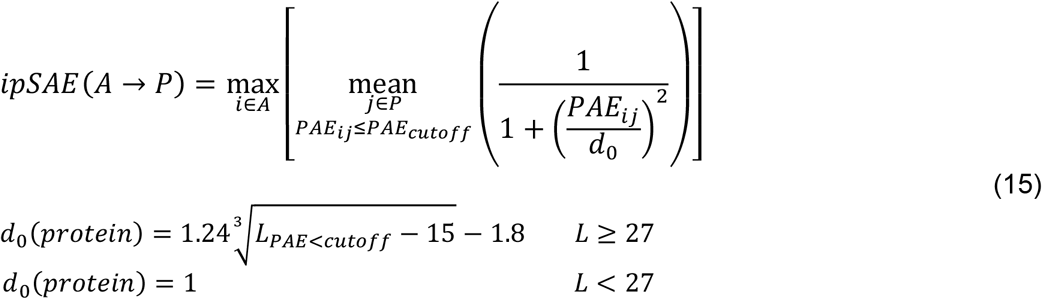

In Eq 15, *L*_*PAE*<*cutoff*_ is the number of unique residues in chain B that have *PAE*_*ij*_ < *cutoff* given the identity of the aligned residue *i*. In the current github code, we use a minimum value of 1 for *d*_0_, since Zhang and Skolnick did not test the fit for proteins shorter than 30 amino acids (*d*_0_=1 for L∼26.5), and the denominator in Eq. 15 starts to blow up for values << 1.0, which may not be realistic or helpful. In the AlphaFold code, the minimum value of *L* is set to 19, since *L*=18 produces a negative number. *L*=19 produces a *d*_0_ of 0.168.

For a given protein-protein chain pair, *P*_1_, *P*_2_, the *ipSAE* score is the maximum of the two asymmetric values:

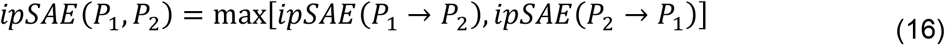

*ipSAE* can be calculated for every pair of chains in a multi-protein complex from the *PAE* matrices from the AlphaFold2 or AlphaFold3 output json files as well as the Boltz-2 npz files.

For protein-ligand complexes, AlphaFold3 and Boltz-2 calculate *ipTM* from the probability distribution over the *PAE* values in both directions. So, from the two asymmetric values for the protein (*P*) and ligand (*L*), the *ipTM* is calculated from their maximum value:

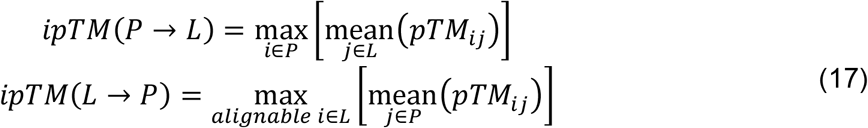

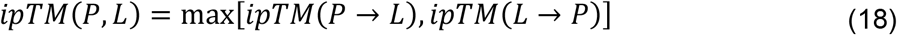

In line 1 of Eq. 17, for every token *i* of a protein, AF3 and Boltz-2 calculate the average of the *pTM* expression over the tokens *j* of the ligand, i.e. all heavy atoms. The asymmetric value comes from the maximum over the tokens in *P*, which consist of one token for each standard amino acid and one token per heavy atom in non-standard or modified amino acids. In Line 2 of Eq 17, for every *alignable* token of the ligand, AF3 and Boltz-2 calculate the average of the TM expression over the protein residues, with the resulting value from the maximum taken over the ligand token atoms. Both programs calculate *PAE*s aligned on a ligand token by taking two nearby atoms in the ligand to create an orientable reference frame. As noted earlier, for some molecules, like an ion or a linear molecule (e.g. carbon monoxide), it is not possible to create an orientable reference frame, and these tokens do not participate in the *ipTM* expressions as alignable tokens (although they can be scored tokens).

For *ipTM*, the same value of *d*_0_ is used in both directions. In AF3, the value of *d*_0_ is calculated as the sum of the number of tokens in the protein and the number of heavy atoms (tokens) in the ligand. In AlphaFold3, a modified amino acid has one token per heavy atom, which are all counted in the sum of tokens for calculating *d*_0_. In Boltz-2, the value of *d*_0_ is calculated from the number of tokens in the entire system, including all proteins and ligands.

This formulation is problematic in two ways. First, for structure prediction of protein-ligand complexes, it would generally be desirable to score the whole ligand. However, when *ipTM* arises from *ipTM*(*L* → *P*), only one ligand frame contributes to scoring the protein residues. Second, when aligning on a protein residue, the *d*_0_ implies scoring an object that is the size of a protein with the number of tokens in the binding protein plus the number of tokens in the ligand. However, a ligand typically has 10s of atoms, not 100s of protein residues. Thus, the *PAE* values are scaled downwards by a very large *d*_0_. This introduces an artifact such that the size of the protein construct affects the *ipTM* of the ligand, which is not realistic or useful. Increasing the number of tokens used to calculate *d*_0_ raises the TM-score contribution of any fixed *PAE* value, potentially inflating *ipTM* independently of improvements in the modeled interface.

To fix these problems, we focus on the asymmetric P→L score, aligning on protein residues, scoring the whole ligand without a *PAE* cutoff (since disordered residues and irrelevant protein domains are not generally a factor for binding a ligand), and setting *d*_0_ to a low, constant value.

Currently, we set this to *d*_0_ = 2.0, which we justify in the Results, although more extensive benchmarking in the future may be used to optimize this value. Thus, we define for a molecule A which is either a protein or a nucleic acid interacting with a ligand *L*:

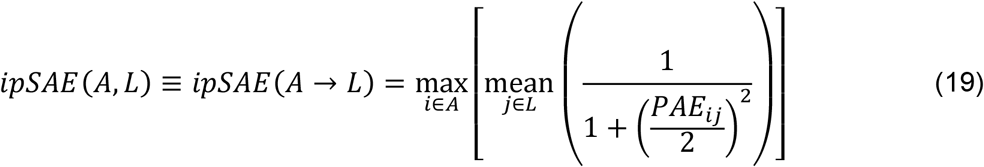

We also calculate *ipSAE*(*L* → *A*), where we align on ligand tokens and score the tokens in molecule A. It may serve some purposes as described in the Results. For ligand-ligand interactions, it makes sense to take the maximum of the two asymmetric values, so

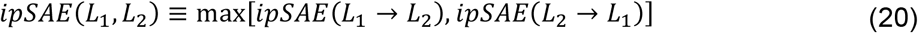

Finally, we investigate how to score nucleic acids (see Results): whether there is a *PAE* cutoff and how *d*_0_ should be determined. For large nucleic acids which contain nucleotides far away from the protein binding site, their *PAE* values may increase due to flexibility in the nucleic acid. To account for this, it makes sense to include a *PAE* cutoff, just as we do for scoring proteins. However, since the scored molecule is not a protein but instead usually involves a small number of nucleic acid bases, we set *d*_0_ to a constant. Currently, we use a value of 2.0, which benchmarking shows to be reasonable. So, for the interaction of a protein *P* and a nucleic acid *N*:

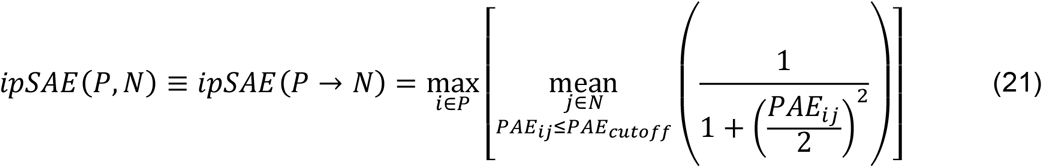

However, when both chains are nucleic acids, we use the max of the two asymmetric values, with

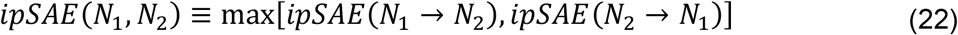

We can summarize the rules for the *ipSAE* score:

1. For all molecule pairs (except those with atomic ions), we calculate and output the two asymmetric scores, *ipSAE*(*A* → *B*) and *ipSAE*(*B* → *A*). It is not possible to align on ions, so we only provide the asymmetric score, *ipSAE*(*A* → *Ion*) and the other is “NA”.
2. When proteins are scored, we use a *PAE* cutoff (typically 10 Å) to eliminate the effect of unbound disordered regions and domains in the scored chain that are not part of the evolved interface. We base *d*_0_ on the number of residues in the scored chain that pass the cutoff.
3. When nucleic acids are scored, we also use a *PAE* cutoff (typically 10 Å) to eliminate the parts of the nucleic acid chain that are not part of the evolved interface (e.g., in long RNA molecules or DNA duplexes with regions distant from the protein-binding site). We set *d*_0_ to a constant (currently 2.0).
4. When ligands or atomic ions are scored, we do not use a *PAE* cutoff and we set *d*_0_ to a constant (currently 2.0).
5. For molecule pairs of the same type (except ions), since there is no physical distinction between the two directions, we define the *ipSAE* score as the maximum of the two asymmetric values. Since it is not possible to align on ions, there is no *ipSAE* score for ion-ion pairs.
6. For dissimilar molecule pairs, *ipSAE* is the asymmetric value such that the aligned tokens are in the more complex molecule and the scored tokens are in the less complex molecule, such that protein > nucleic acid > ligand > ion.

### Ligand structure prediction generation and benchmark evaluation

Protein–ligand complexes were selected from the FoldBench benchmark (13). We retained assemblies containing one protein polymer chain and one noncovalently bound small-molecule ligand, with no additional polymer chains and no more than four monatomic ions. Protein lengths were restricted to 80–800 residues and ligand sizes to 8–70 heavy atoms. Common crystallization additives were excluded, and each ligand Chemical Component Dictionary (CCD) identifier was represented only once. The resulting 102 complexes were ordered by ligand heavy-atom count, and 20 approximately evenly distributed targets were selected as a development set (Dev20). The remaining 82 targets comprised the test set (Test82).

Structure predictions were performed with AlphaFold3 (Git commit c0f97eda2f1f482fd94d3a38bece18c7069b4a5c, August 19, 2026), using a locally built Docker image dated September 2, 2026, and Boltz-2 (version 2.2.1). Input structures contained the protein sequence, ligand specified by its CCD identifier, and retained ions where applicable. For each target and prediction program, five random seeds (42–46) were used, with five models generated per seed (samples 0–4), producing 25 models per target. Thus, Dev20 contained 500 models and Test82 contained 2,050 models for each program, for a total of 5,100 predicted structures.

Ligand RMSDs and protein–ligand interface accuracy metrics were calculated using OpenStructure (version 2.8.0) with the compare-ligand-structures procedure and the --lddt-pli and --rmsd options. Ligand RMSD was calculated using the binding-site-superposed, symmetry-corrected RMSD (BiSyRMSD) method. The protein binding site was defined from residues having at least one atom within 4 Å of the ligand in the experimental structure (the OpenStructure default). Corresponding protein residues in the predicted and experimental binding sites were identified by sequence alignment, and their backbone Cα atoms were used for least-squares superposition. Following this superposition, RMSD was calculated over the ligand heavy atoms. Chemically equivalent ligand atom mappings arising from molecular symmetry were enumerated, and the minimum RMSD was retained. Thus, ligand RMSD measurements account for symmetric atoms and groups and do not depend solely on atom names. OpenStructure also provided the protein– ligand interface lDDT (lDDT-PLi) and binding-pocket lDDT (lDDT-LP) scores.

For the classification benchmarks, models with ligand RMSD ≤ 5 Å were designated positive (approximately correct ligand placement), and those with RMSD > 5 Å were designated negative. We evaluated the protein-to-ligand *ipSAE* score, *ipSAE*(P→L), calculated with a fixed *d*_0_=2 Å and without a *PAE* cutoff, so that all ligand heavy-atom tokens were included in the score. Models were classified as positive when *ipSAE* was greater than or equal to a specified threshold. We calculated sensitivity (true-positive rate, TPR), specificity (true-negative rate, TNR), positive predictive value (PPV), negative predictive value (NPV), and balanced accuracy (BACC), defined as (TPR + TNR)/2. The area under the receiver operating characteristic curve (AUROC) was calculated from the continuous *ipSAE* values over all possible thresholds.

Optimal thresholds were identified separately for each program and dataset by evaluating the observed *ipSAE* score boundaries and selecting those that maximized BACC. Where multiple thresholds yielded identical maximum BACC, simpler reference values were preferred when applicable. The development set was used to select fixed *d*_0_ and reference thresholds of 0.5 for AF3 and 0.8 for Boltz-2. These reference thresholds were subsequently evaluated on Test82 without further adjustment. Optimal thresholds calculated independently on Test82 are reported for comparison but were not used to select the reference thresholds. Because the 25 models generated for each target are not independent biological examples, the resulting classification statistics describe performance across predicted models and should not be interpreted as estimates based on 2,050 independent targets.

### Protein–DNA and protein–RNA benchmark evaluation

Protein–DNA and protein–RNA complexes were selected from the FoldBench benchmark (13). Predictions were evaluated at the level of individual protein–nucleic acid interfaces, since a single molecular assembly may contain more than one such interface. The final protein–DNA benchmark comprised 158 interfaces from 96 assemblies for AlphaFold3 and 144 interfaces from 90 assemblies for Boltz-2. For protein–RNA complexes, we used a smaller preliminary subset selected from assemblies for which reference structures and prediction-evaluation data were available, with the aim of including a range of structural accuracies. This subset comprised 27 interfaces from 20 assemblies for AlphaFold3 and 19 interfaces from 15 assemblies for Boltz-2. Five RNA assemblies could not be completed with Boltz-2 because of their large size. Predictions were generated using five random seeds (42–46), with five models per seed, yielding 25 models per assembly. After matching predicted structures to their corresponding experimental interfaces and retaining valid evaluations, the datasets comprised 3,950 and 3,600 protein–DNA interface predictions for AlphaFold3 and Boltz-2, respectively, and 658 and 456 protein–RNA interface predictions.

Structural accuracy was assessed using DockQv2 as implemented in the FoldBench evaluation code. DockQv2 thresholds of 0.23, 0.49, and 0.80 were used to classify models as having acceptable-or-better, medium-or-better, and high structural accuracy, respectively. Each threshold was evaluated independently, with models above or equal to the threshold classified as positive and models below the threshold classified as negative.

For each protein–nucleic acid interface, we calculated directional *ipSAE* scores using the protein as the alignment molecule and the nucleic acid as the scored molecule (P→N), as well as the reverse direction (N→P). When scoring nucleic acids, we included only nucleotides with *PAE* ≤ 10 Å for each aligned protein token. We evaluated constant values of *d*_0_ = 1, 2, 3, 4, and 5 Å, as well as the length-dependent *d*_0_ obtained from the TM-score equation (Eq. 2), with *L* equal to the combined numbers of protein and nucleic acid tokens. For the reverse direction (N→P), we used a *PAE* cutoff of 10 Å for scored protein residues and calculated *d*_0_ from the number of residues passing the cutoff, as in the original *ipSAE* formulation. The native *ipTM* scores from AlphaFold3 and Boltz-2 were retained for comparison.

The ability of each score to discriminate accurate from inaccurate models was evaluated by calculating sensitivity (TPR), specificity (TNR), and balanced accuracy (BACC = (TPR + TNR)/2) at different score thresholds, with equal weighting of interfaces. For each *d*_0_ value and DockQv2 accuracy threshold, we determined the maximum BACC over possible *ipSAE* classification thresholds. We also evaluated fixed *ipSAE*(P→N) thresholds of 0.5 for AlphaFold3 and 0.8 for Boltz-2, corresponding to the reference thresholds selected from the protein–ligand development benchmark. These thresholds were not separately optimized on the DNA or RNA datasets.

## RESULTS

### ipSAE *for ligands and atomic ions*

To explore the choices for an *ipSAE* score for ligands, we chose targets from the FoldBench benchmark (13). FoldBench is a broad, low-homology benchmark for all-atom biomolecular structure prediction. The current protein–ligand subset contains 558 protein–ligand interfaces from PDB structures deposited after the AF3 validation cutoff (January 13, 2023) and before November 1, 2024. For protein–ligand interfaces, FoldBench excluded a target when the protein had >40% sequence identity to a protein in an AF3 training complex and the ligand had >50% Tanimoto similarity to the ligand in that same complex. They also randomly retained one interface per ligand and applied additional ligand-quality filters.

We further filtered the FoldBench set as described in Methods, keeping only those available to AF3 and Boltz-2, resulting in 102 structures. We sorted the candidates by ligand heavy-atom count and selected 20 approximately evenly spread across the ligand-size range as a *development set* (Dev20_set) and used the remaining 82 as a *held-out test set* (Test82_set). We ran AlphaFold3 and Boltz-2 with seeds 42-46, producing 25 models per target. The sets are listed in the **Appendix**.

For both programs, we ran the Dev20_set first and calculated *ipSAE*(P→L), aligning on protein residues and scoring the ligand, at different fixed *d*_0_ values (1,2,3,4,5) as well as the AlphaFold3 default, which uses the *d*_0_ formula from the TM score paper (Eq. 2) where *L* is the number of residues in the protein plus the number of tokens (heavy atoms) in the ligand. The results are shown in **Figure 3**, where we plot lDDT-PLi, the protein-ligand local Distance Difference Test values (14) measured by FoldBench, vs the different possible *ipSAE*(P→L) scores. In **Figure 3**, the points are colored by whether they have approximately the right orientation in the right pocket (blue, RMSD ≤ 2.0 Å), an intermediate accuracy (orange, 2.0 < RMSD ≤ 5 Å), or an incorrect pose (green, RMSD > 5 Å). For most targets, the 25 predictions gave consistently accurate or inaccurate ligand placements, with relatively few targets producing mixtures of models on opposite sides of the 5 Å RMSD threshold. Since the blue and orange points are all above lDDT-PLi of 0.7 and the green points are all below lDDT-PLi of 0.5, we use 5.0 Å as a cutoff to define positive predictions (RMSD ≤ 5.0 Å) and negative predictions (RMSD>5.0 Å).

**Figure 3.**
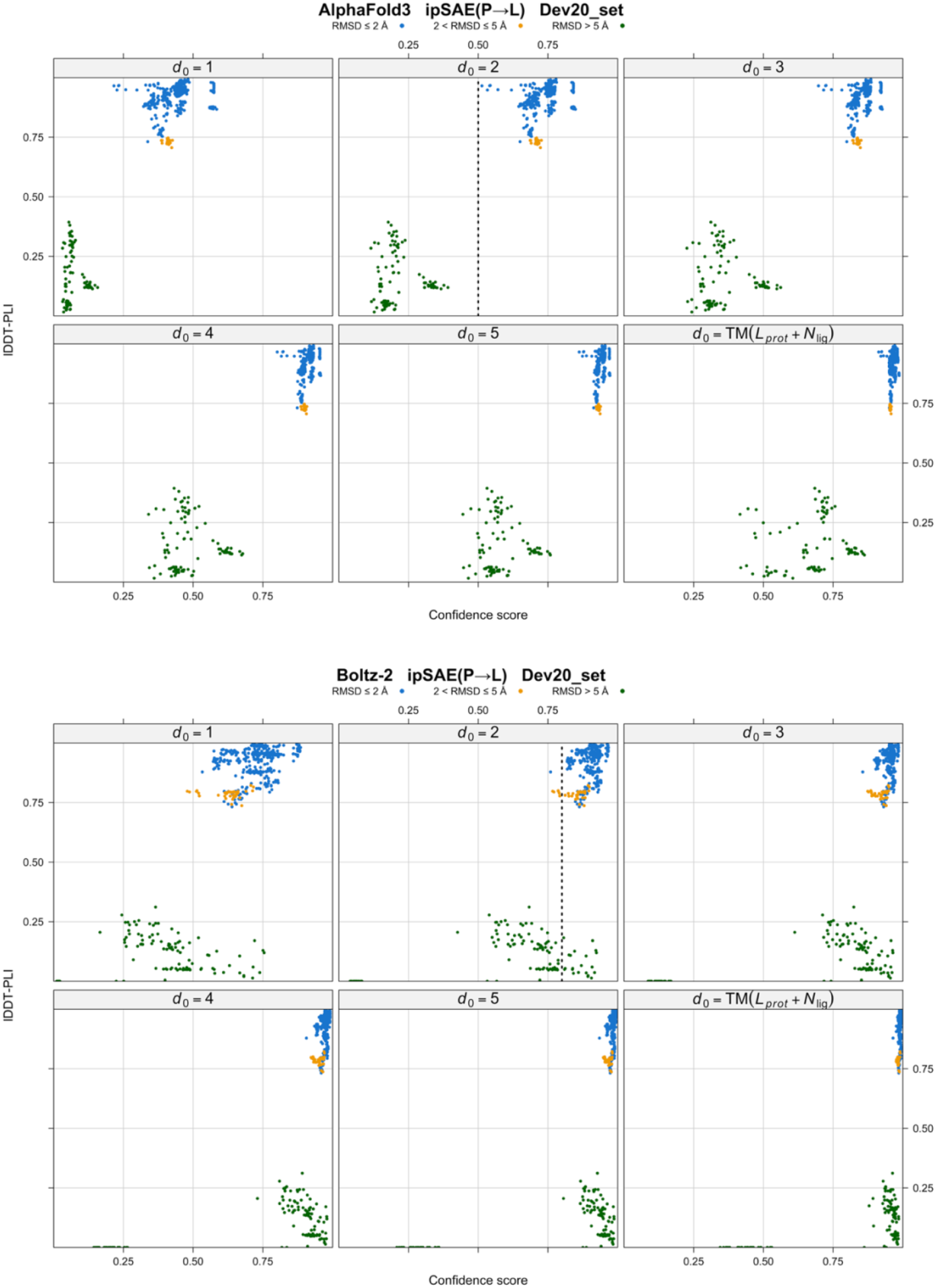
Comparison of various implementations of the *d*_0_ parameter for the asymmetric *ipSAE*(P→L) score for ligands in the development set of 20 ligands. Results are shown for 25 models with AlphaFold3 (top set of figures) and Boltz-2 (bottom set of figures) for each of 20 targets. *d*_0_ was either set as a constant with values of 1, 2, 3, 4, and 5 (top 3 panels and first 2 panels of bottom row), or set to the value prescribed by the TM formula (Eq. 2) with *L* given by the number of tokens in the protein plus the number of tokens in the ligand (bottom right). For AlphaFold3, the *d*_0_ = 2 panel shows the widest dispersion of scores and the largest separation of mean score for well modeled structures (*lDDT-PLi* > 0.7) and poorly modeled structures (*lDDT-PLi* < 0.4). For Boltz-2, the *d*_0_ = 1 panel shows the widest dispersion of scores. However, the *d*_0_ = 2 panel also shows a significant separation of positive and negative models at a score cutoff of 0.8.

Given the RMSD distributions per target, correlation coefficients of accuracy vs score for one target were not very meaningful. The goal is a score that can meaningfully separate targets for which all or most models are positive predictions and those for which few or none are. For AF3, the largest difference of average scores for modeled structures better or worse than 5 Å RMSD is largest for *d*_0_=2 (0.74 and 0.21 respectively, a difference of 0.53), and *ipSAE* of 0.5 completely separates the two sets of models. By contrast, the AlphaFold3 *d*_0_ default from the TM formula (bottom right) has corresponding values of 0.97, 0.68 and a difference of 0.29. A value of *d*_0_=3 is similar to *d*_0_=2 with averages of 0.86 and 0.37, and a difference of 0.49. Thus, *d*_0_=2 seems to be a reasonable choice for *ipSAE*(P→L) for AlphaFold3.

Although *d*_0_=1 also provides useful discrimination, particularly for Boltz-2, *d*_0_=2 produces a greater separation of mean scores for accurate and inaccurate AF3 models and retains useful discrimination for Boltz-2. We therefore selected *d*_0_=2 as a common value for both programs. The resulting score thresholds differ between AF3 and Boltz-2, reflecting differences in their *PAE* distributions. For Boltz-2, a value of *d*_0_=1 gives the largest difference in positive and negative scores and a potential score cutoff of 0.5. However, a value of *d*_0_=2 works equally well for separating positive and negative predictions with a score cutoff of 0.8.

Next, we examined the distribution of scores for *ipSAE*(L→P) and the native *ipTM* scores vs lDDT-PLi for the Dev20_set for each program. These are shown with *ipSAE*(P→L) with *d*_0_=2 in **Figure 4** (AF3 top row, Boltz-2 bottom row). The *ipSAE*(L→P) (center panels in each row) have wider dispersions of scores than *ipSAE*(P→L) (left panels in each row) for both accurately and inaccurately predicted structures, and there is more significant overlap in score between the two populations compared to *ipSAE*(P→L), especially for Boltz-2. *ipTM* pushes all the scores to higher values than *ipSAE*(P→L) and all are above 0.5. For Boltz-2, the *ipTM* scores of incorrect models are almost all above 0.9 and overlap significantly with the *ipTM* scores for correct models.

**Figure 4.**
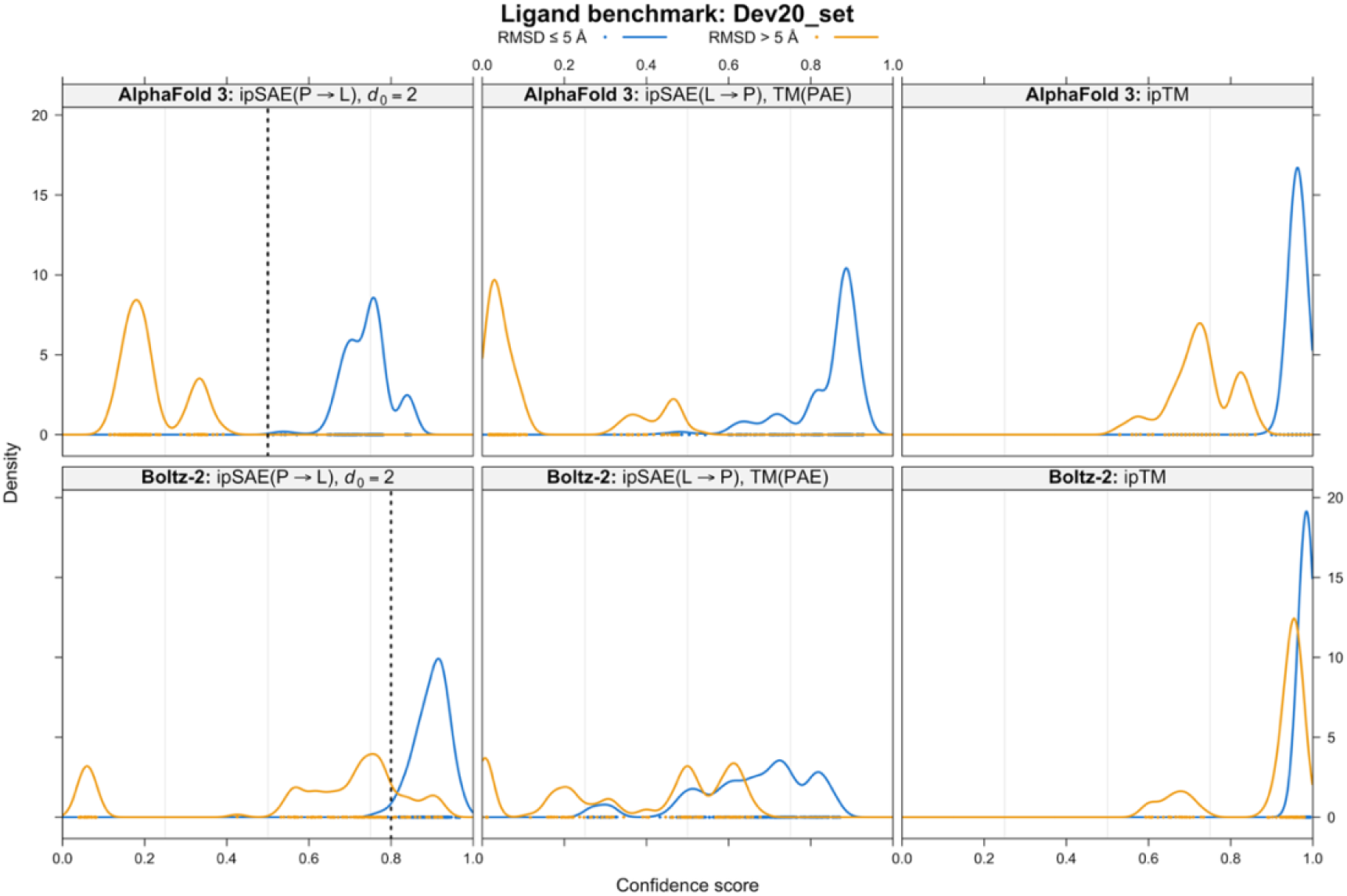
Kernel density estimates of *ipSAE*(P→L) scores with *d*_0_=2 (left panels) with *ipSAE*(L→P) scores (center panels), and native *ipTM* scores (right panel) for AlphaFold3 (top row) and Boltz-2 (bottom row) for the Dev20_set. Models with RMSD ≤ 5 Å are in blue points and curves and those with RMSD > 5 are in orange. For AlphaFold3, some of the *ipSAE*(L→P) values are larger than the *ipSAE*(P→L) values, due to the *d*_0_ normalization constant from the number of protein residues that pass the *PAE* cutoff. For Boltz-2, the L→P values are lower than the P→L values. For Boltz-2, the *ipTM* values for models with RMSD ≤ 5 Å are at very high score values and overlap the values for models with RMSD > 5 Å.

Finally, we examined whether adding a *PAE* cutoff affects the score. The scores with *PAE* cutoff were identical to those without cutoff (not shown). We choose not to include a *PAE* cutoff for *ipSAE* for ligands, so that entire ligands are scored.

From **Figures 3** and **4**, we tentatively conclude that for ligands, *ipSAE* should be defined as the asymmetric score *ipSAE*(P→L) with *d*_0_=2 and no *PAE* cutoff. To determine if this is satisfactory, we processed the same data for the Test82_set; the results are shown in **Figures 5 and 6**.

**Figure 5.**
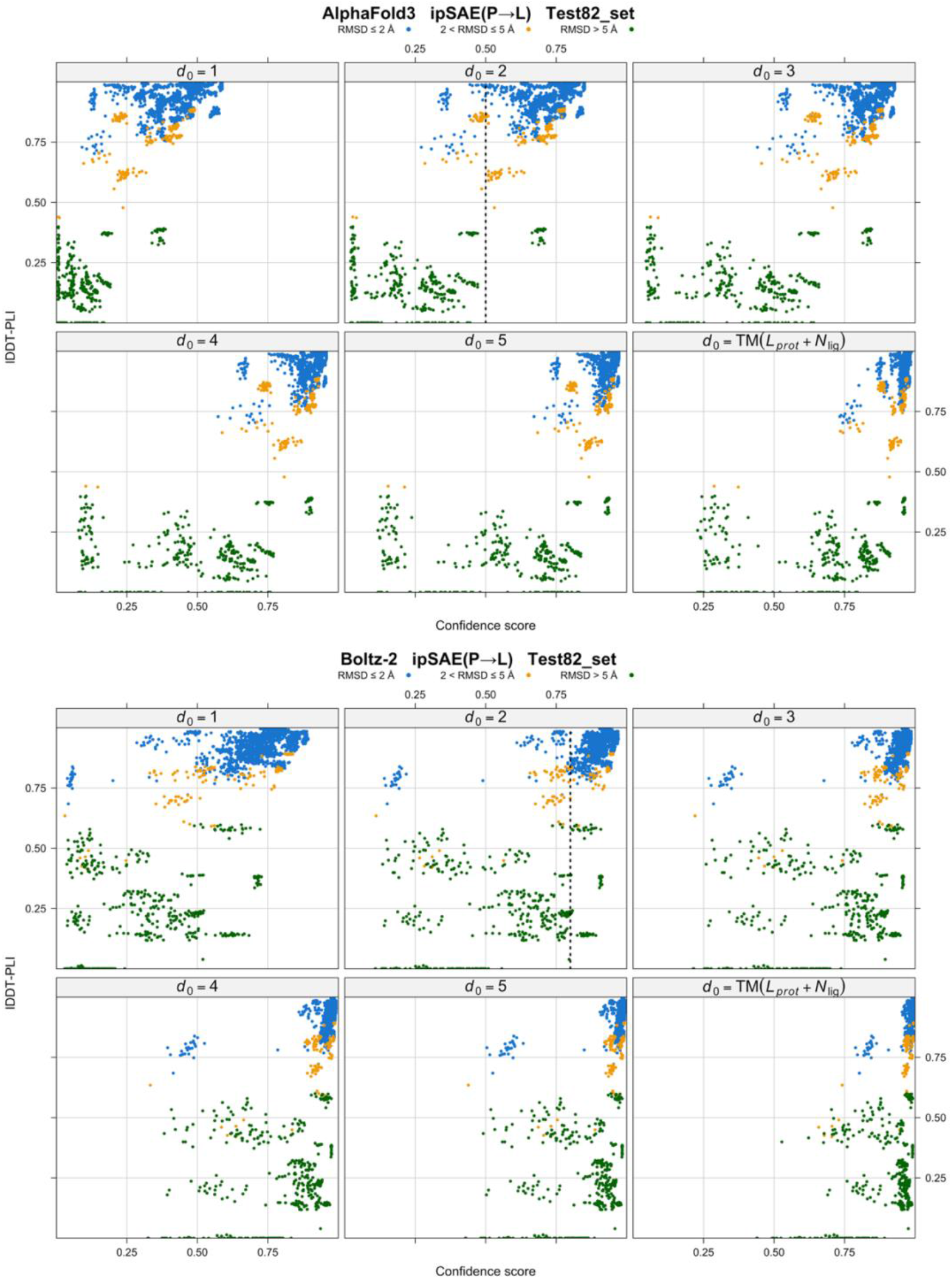
Comparison of various implementations of the *d*_0_ parameter for the asymmetric *ipSAE*(P→L) score for ligands in the testing set of 82 ligands from FoldBench. Results are shown for 25 models with AlphaFold3 (top set of figures) and Boltz-2 (bottom set of figures) for each of 82 targets. *d*_0_ was either set as a constant with values of 1, 2, 3, 4, and 5 (top 3 panels and first 2 panels of bottom row), or set to the value prescribed by the TM formula (Eq. 2) with *L* given by the number of tokens in the protein plus the number of tokens in the ligand (bottom right). For AlphaFold3, we show a cutoff of 0.5 for approximately correct and incorrect models for *d*_0_=2. For Boltz-2, we show a cutoff of 0.8 for approximately correct and incorrect models for *d*_0_=2.

**Figure 6.**
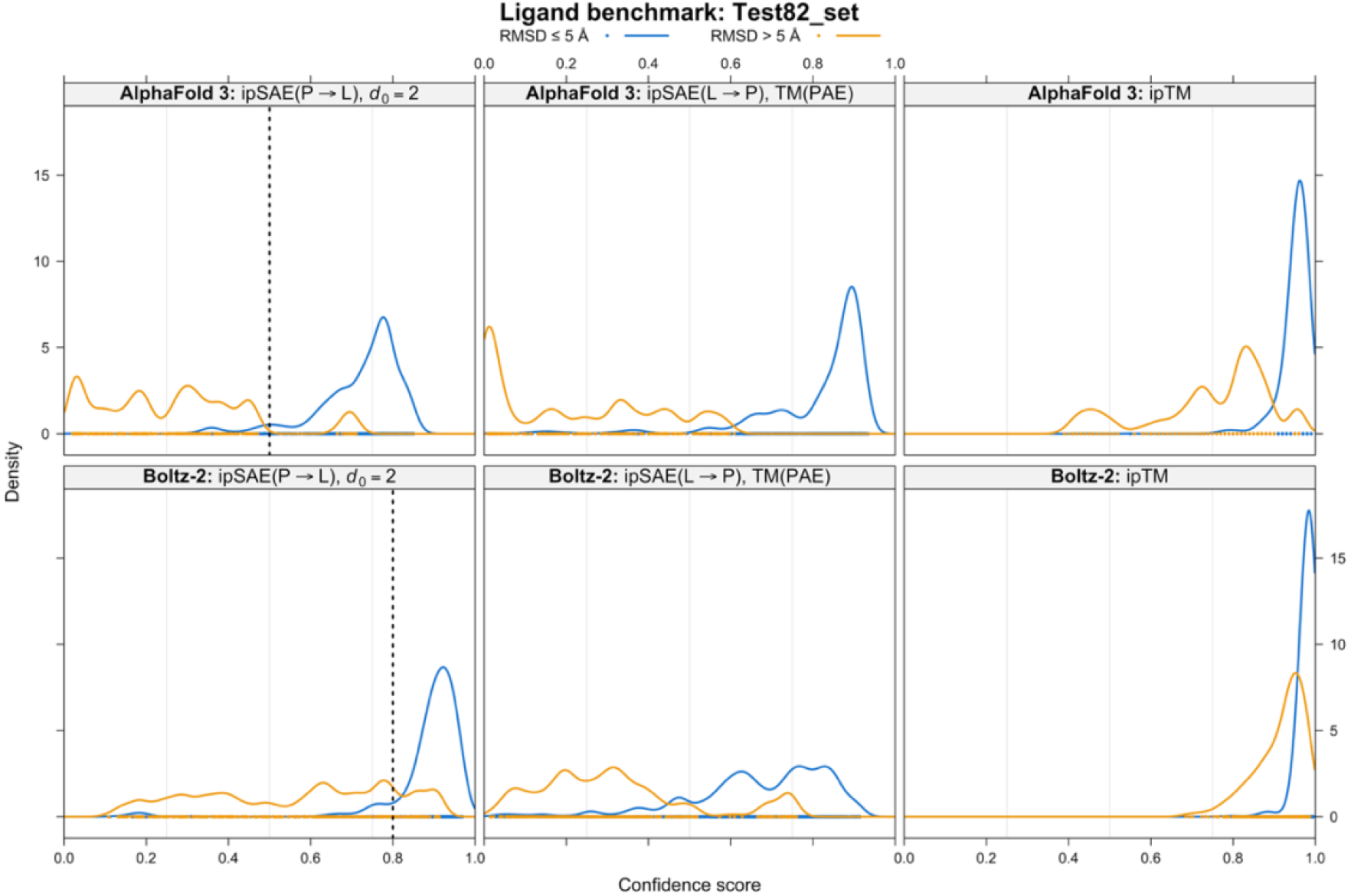
Kernel density estimate comparison of *ipSAE*(P→L) with *d*_0_=2 (left panels) with *ipSAE*(L→P) (center panels), and native *ipTM* (right panel) for AlphaFold3 (top row) and Boltz-2 (bottom row) for the Test82_set.

**Figures 5** and **6** show similar properties of the different scores and the same distinct features of the two programs. At *d*_0_=2, AF3 has a relatively clean cutoff of *ipSAE* = 0.5 between accurate and inaccurate models. The models with *ipSAE*(P→L) > 0.65 and RMSD>5 all come from one target, PDB 8cey. This is a long ligand that AF3 flips over end-to-end, so the ligand is in the right pocket but in a reversed orientation. For Boltz-2 at *d*_0_=2, a cutoff of *ipSAE* = 0.8 performs the same function (**Figure 5**). Almost all the models with Boltz-2 *ipSAE*(P→L) > 0.8 and RMSD>5 Å come from three targets (PDB 8cey, 8ouv, 8pqg). The native *ipTM* scores (**Figure 6, lower right**) push all the values to very high values and higher overlap between positive and negative models than *ipSAE*(P→L).

To assess the cutoffs more quantitatively, in **Table 1**, we show the optimal cutoffs for balanced accuracy (BACC=(TPR+TNR)/2), TPR, TNR, and AUROC for the full set of 102 ligands. Positive models are defined as those with RMSD ≤ 5 Å (most of which are below 2 Å, blue dots in **Figure 5**) and negative models with RMSD > 5 Å. The table provides the same values for rounded-value reference cutoffs of 0.5 and 0.8 for AlphaFold3 and Boltz-2 respectively. For AF3, the optimal cutoff is 0.471, justifying the 0.5 cutoff as a rounded value. For Boltz-2, the optimal cutoff is 0.812, resulting in a rounded value of 0.8. The classification performance changes very little between the optimal cutoff and the reference cutoff values. The final reference BACC values for the full set of 102 targets are 0.957 and 0.876 for AlphaFold3 and Boltz-2 respectively, versus 0.951 and 0.871 for the optimal values. The *ipTM* values are 0.945 and 0.879. While the AF3 value for *ipSAE* is slightly higher for AF3, it is very slightly lower for Boltz-2. However, the optimal cutoffs for *ipTM* are very high, 0.910 and 0.969, indicating the challenge of setting a cutoff for *ipTM* for discrimination of accurate and inaccurate models. The *ipSAE* scores present a much larger dynamic range, which is useful in a discriminative score.

**Table 1.**
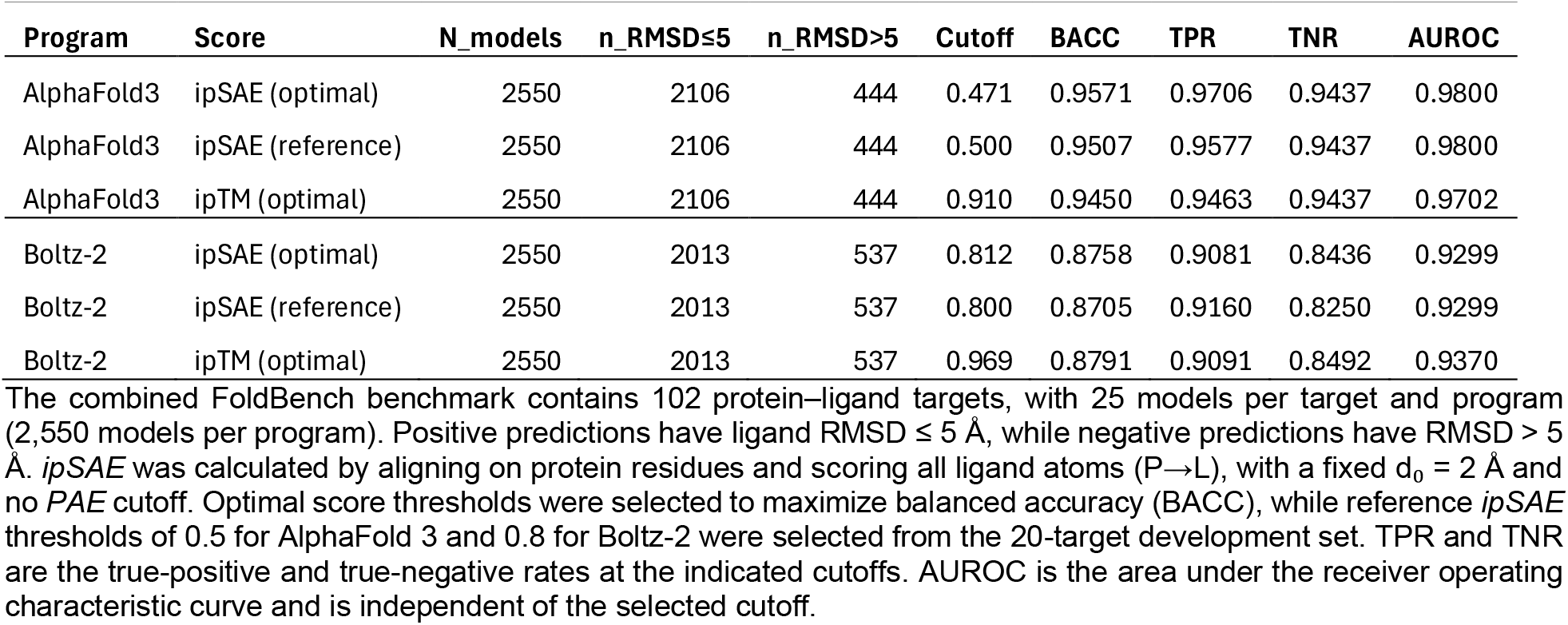
Comparison of *ipSAE* and native *ipTM* for classification of protein–ligand complex predictions from AlphaFold 3 and Boltz-2.

### ipSAE *for protein-ligand interactions: Test case AURKA with cognate and non-cognate ligands*

For *ipTM*, both AlphaFold3 and Boltz-2 align on protein residues and score the ligand and align on ligand tokens and score the protein. In the latter case, it is possible that a fragment of a ligand may bind quite well to a site, but some other part of a ligand does not. For instance, an ATP binding site may accommodate NAD or FAD at the adenosine, but the protein may not accommodate the flavin or nicotinamide group. By contrast, the defined *ipSAE* score for ligands aligns only on the protein residue and scores the entire ligand with a constant value of *d*_0_. To test the *ipSAE* metric, we ran AlphaFold3 and Boltz-2 structure predictions on Aurora A kinase (AURKA) with a variety of true ligands and decoy ligands and with a fragment of the activator protein TPX2 (residues 1-43). The structure of AURKA with ADP, phosphorylated activation loop (on Thr287 and Thr288), and TPX2 is used as a basis for these calculations (PDB: 1OL5 (15)).

The results in **Figure 7** show that while *ipSAE* and *ipTM* rank the compounds in much the same order, *ipSAE* has much greater dynamic range than *ipTM* for both AlphaFold3 (top left panel) and Boltz-2 (bottom left panel). The *ipSAE* ranges for AF3 and Boltz-2 are [0.231, 0.766] and [0.061, 0.915] respectively, while the *ipTM* ranges are [0.810, 0.970] and [0.676, 0.987]. For both programs, *ipTM* is higher than *ipSAE* for all ligands. This is primarily because for *ipTM*, the *PAE* matrix is scaled by a *d*_0_ parameter calculated from the length of the protein plus the number of ligand tokens. For AURKA and a ligand with 30 heavy atoms (like ATP), *d*_0_ is 6.5 Å in AlphaFold3. Because Boltz-2 bases *d*_0_ on the total number of tokens in the system, the number of residues in TPX2 and the two Mg^2+^ ions increase the *d*_0_ parameter to 6.88 Å. *ipSAE* uses a value of 2.0. Although ATP, ADP, and possibly AMP are the most biologically relevant ligands in this set, the other simple nucleotide phosphates (GDP, GTP, UTP, CTP) also score reasonably well with *ipSAE* with values ranging from about 0.6 to 0.8 in AF3 and 0.8 to 0.9 in Boltz-2. These values are consistent with the cutoff values we chose for AF3 and Boltz-2 of 0.5 and 0.8 respectively. We should remember that the *PAE* matrix reflects predicted structural accuracy and not binding affinity. The kinase active site has evolved to bind a nucleotide base, ribose, and the phosphate segment. The ligands most distinct from nucleotides (OLA, PLM, HEM) score much more poorly with *ipSAE* than *ipTM* for both programs, because of the increased dynamic range.

**Figure 7.**
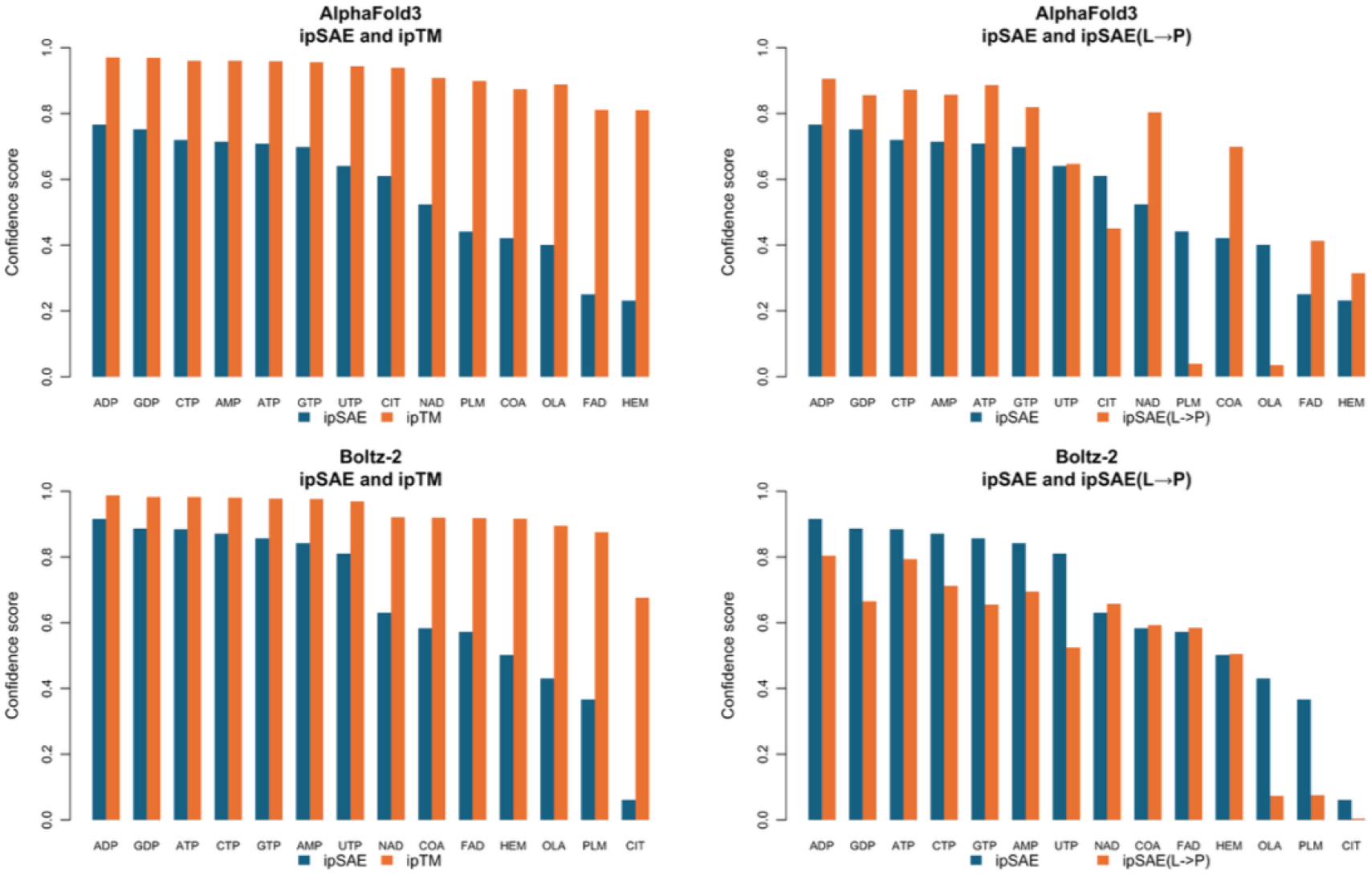
*ipSAE* and *ipTM* scores for ligand complexes of AURKA kinase domain and a fragment of TPX2. For the canonical nucleotides (ATP, ADP, GDP, CTP, AMP, GTP, UTP), we included two magnesium ions. Both Thr287 and Thr288 are phosphorylated. CIT is citrate, PLM is palmitic acid, OLA is oleic acid, HEM is heme. A total of 25 models were produced for each target complex with each program. Left column: comparison of *ipSAE* and *ipTM* for AlphaFold3 and Boltz-2 modeling of natural and decoy ligands. While the ranking of ligands is very similar, *ipSAE* provides substantially greater dynamic range than *ipTM*, particularly by assigning lower confidence scores to the noncognate ligands CIT, PLM, OLA, and HEM. *ipSAE* aligns on protein residues and scores all ligand atoms. We can also calculate the reverse score, aligning on ligand tokens and scoring the whole protein (with a *PAE* cutoff of 10.0). For AlphaFold3, the *ipSAE*(L→P) scores (top right) are higher than *ipSAE*, except for three non-biological ligands: CIT, PLM, and OLA. However, adenosine diphosphate-based ligands score very high on *ipSAE*(L→P), because atoms in the adenine ring score the ATP binding site very highly and are responsible for the score. For Boltz-2, *ipSAE* is higher for most ligands, except for NAD, COA, FAD, and HEM.

We also compared the *ipSAE*(L→P) values with *ipSAE*(P→L). For *ipSAE*(L→P), we use a *PAE* cutoff of 10 Å to score the protein and calculate *d*_0_ from the number of residues in the protein that pass the *PAE* cutoff. This is necessary because the protein may contain large regions that are not part of the ligand-binding domain and these would inflate *d*_0_ unnecessarily. For AF3, *ipSAE*(L→P) is larger than *ipSAE* for 11 of the 14 test ligands, with the exceptions of the most non-cognate ligands CIT, OLA, and PLM. Of these 11 ligands, 10 contain nucleotide-like moieties; HEM is the sole exception. Among the nucleotide phosphate ligands, NAD, COA, and FAD have a much higher ratio of *ipSAE*(L→P) to *ipSAE* than the simpler nucleotide phosphates (AMP, ADP, ATP, GTP, UTP, CTP). This is because the *ipSAE* values for NAD, COA, and FAD are lower than the simpler ligands, while the *ipSAE*(L→P) values are comparable. NAD for instance has an *ipSAE*(L→P) value close to that of GTP (0.80 vs 0.82 respectively), while the *ipSAE* values are 0.52 and 0.70 respectively. For both sets of ligands, the *ipSAE*(L→P) values almost all come from aligning on a token in the nucleotide base and scoring the protein, indicating that the network can confidently place the base in the kinase active site. The *ipSAE* values are lower for COA, NAD, and FAD because the entire ligand is scored for each protein residue, so that while the adenine scores well, the non-cognate moiety has poorer *PAE*s with the protein.

Boltz-2 differs from AF3 by having larger *ipSAE* values than *ipSAE*(L→P) for the seven simpler nucleotide phosphates, while for COA, NAD, and FAD, the *ipSAE*(L→P) are slightly larger than *ipSAE*. Again, the *ipSAE*(L→P) values are comparable between the simpler and longer nucleotide phosphates, while the *ipSAE* values are lower for the longer ligands. For the non-cognate ligands, CIT, OLA, and PLM, the *ipSAE*(L→P) values are very small, indicating that no token in the ligand has a clear relationship to residues in the protein.

In sum, the results motivate the rationale that the more useful metric in most instances is *ipSAE* (i.e., *ipSAE*(P→L) with *d*_0_=2 and no *PAE* cutoff) rather than the maximum of the P→L and L→P directions, as in *ipTM*. The L→P values can come from single well-placed tokens in the protein binding site, while the P→L values in *ipSAE* come from scoring the entire ligand.

### ipTM *and* ipSAE *for protein-ligand interactions: A test case with different protein construct lengths*

*ipTM* for ligands is dependent on the protein construct length, which may lead to artifacts when correlating the score to RMSD values or to raw *PAE* values. We explored this with the ligand D-myo-inositol-1,4,5-trisphosphate (commonly abbreviated Ins(1,4,5)P_3_) bound to rat phospholipase C delta 1 (PDB: 1MAI, ligand CCD code: I3P). We tested three different constructs of the human protein: 1) the PH domain alone, residues 1-135; 2) the PH domain with the adjacent EF-hand region (residues 1-300); 3) the full-length protein, which contains the PH domain, the two EF-hand domains, a catalytic X/Y region forming an α/β (TIM) barrel, and a C2 domain (residues 1-756).

For each construct length of human PLCD1, we produced 25 models each with AF3 and Boltz-2 and aligned the PH domain of each model to the structure in PDB 1MAI of the rat PLCD1 PH domain. We then measured the RMSD of the ligand in each model to the ligand in 1MAI. Unexpectedly, the protein-ligand *PAE* matrix and the RMSD of the ligand to the native structure can be affected by the length of the construct (**Table 2, Figure 8**). The reasons for this are not clear but the network seems to be noisier with the addition of extra sequence along with the ligand binding PH domain. In any case, the *ipSAE* values for AF3 are 0.75, 0.59, 0.33 across the constructs from short to long and for Boltz-2 the values are 0.81, 0.75, and 0.19. The *ipTM* values fail to capture the increased uncertainty in the structure predictions, reflected by the average *PAE* values, and the decrease in structure prediction accuracy, reflected by the increasing RMSD. The AF3 values are (0.94, 0.93, 0.91) and the Boltz-2 values are (0.94, 0.96, 0.88). The Boltz-2 value actually goes up for the middle-length vs shortest construct because of the *d*_0_ effect in *ipTM*.

**Table 2.** *ipSAE, ipTM, PAE*, and Ligand RMSDs for PLCD1 models of various lengths.

| Prog | Construct | Len | Ligand *PAE* mean | Ligand RMSD mean | *ipSAE* mean | *ipTM* mean |
|---|---|---|---|---|---|---|
| *AF3* | 1-135 | 135 | 1.151 | 0.371 | 0.7518 | 0.9400 |
| *AF3* | 1-300 | 300 | 1.701 | 0.521 | 0.5879 | 0.9300 |
| *AF3* | 1-756 | 756 | 2.999 | 0.714 | 0.3263 | 0.9076 |
| *Boltz-2* | 1-135 | 135 | 0.963 | 0.438 | 0.8099 | 0.9445 |
| *Boltz-2* | 1-300 | 300 | 1.167 | 0.580 | 0.7478 | 0.9569 |
| *Boltz-2* | 1-756 | 756 | 4.231 | 1.081 | 0.1887 | 0.8806 |

Each mean is over the 25 models for each construct and program. The ligand *PAE* is the mean *PAE* of ligand atoms when aligning on the protein residue that produces the *ipSAE* score. The ligand RMSD was calculated after superposition of the model’s PH domain to the PH domain in PDB 1MAI.

**Figure 8.**
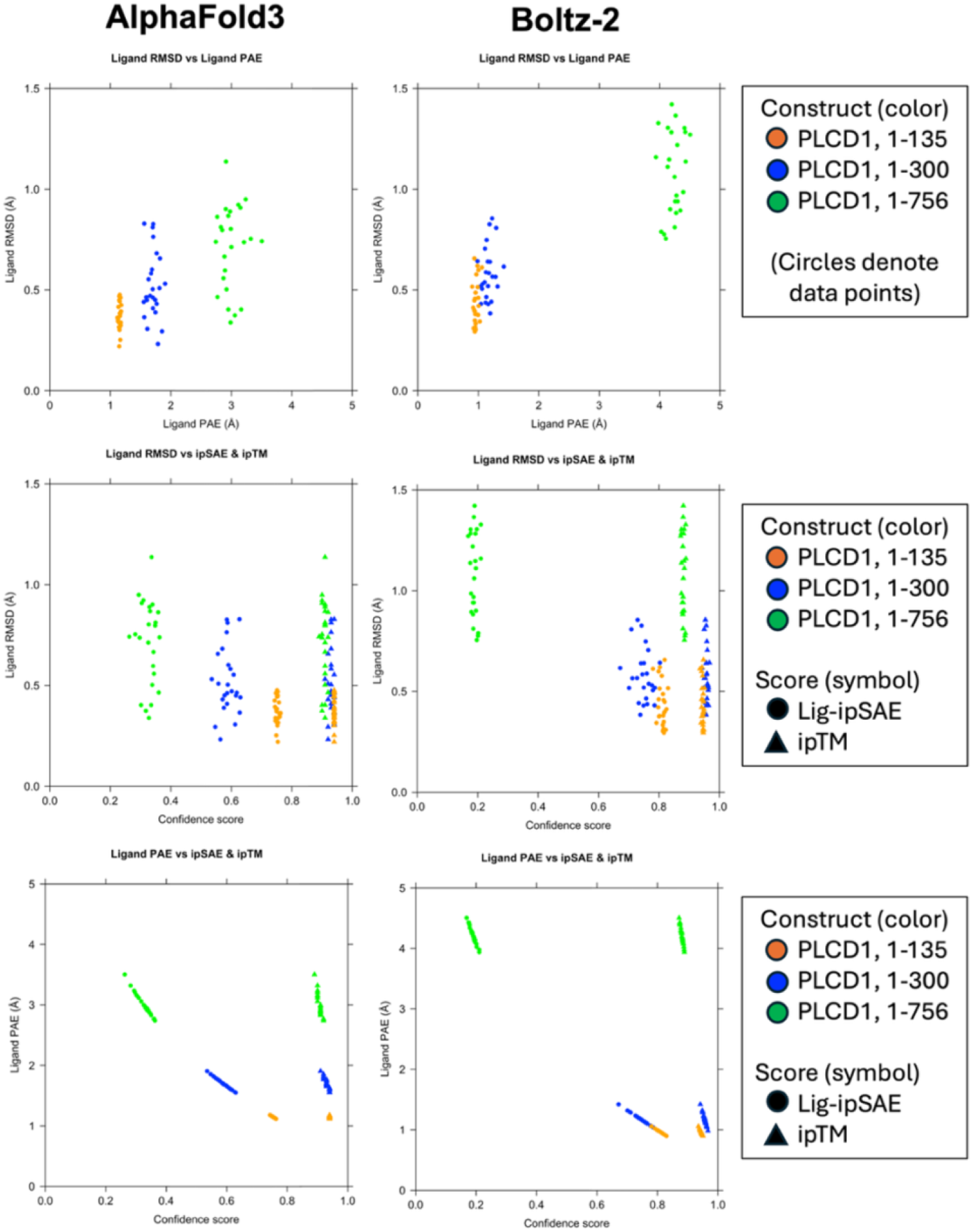
Comparison of *ipSAE* and *ipTM* with *PAE* averages and RMSD for PLCD1. For PLCD1, there are three plots for AF3 (left column) and three plots for Boltz-2 (right column). The colors in each plot indicate the construct of the protein target used: blue, then green, and orange, in order of increasing size. The legends to the right list the constructs. The top row provides the RMSD (y-axis) vs an expression of the *PAE* matrix between the protein and the ligand. For this measure, we calculate the average <u>*PAE*</u> of the ligand atoms when aligning on the protein residue that produces the highest *ipSAE*(P→L) score. The middle row shows the RMSD of the ligand after superposition on the experimental structure vs *ipSAE* (filled circles) and *ipTM* (filled triangles). The bottom row shows the minimum average *PAE* of the ligand vs *ipSAE* and *ipTM*.

Although ligand RMSD increases with construct length, the predicted ligand poses remain highly accurate, with mean RMSDs of approximately 0.7 Å and 1.1 Å for the longest constructs from AF3 and Boltz-2, respectively. Nevertheless, their mean *ipSAE* scores fall below the classification thresholds established from FoldBench. Thus, in these examples, *ipSAE* correctly reflects the increased uncertainty predicted by the networks but underestimates the actual accuracy of the ligand poses. The results demonstrate that *ipSAE* thresholds are not universally applicable and that low predicted confidence does not necessarily imply an inaccurate structure. They also suggest that restricting predictions to the ligand-binding domain, when biologically appropriate, may improve confidence calibration.

### DNA and RNA

#### Protein–DNA complexes

We evaluated directional *ipSAE* scores for protein–DNA complexes predicted by AlphaFold3 and Boltz-2, using 3,950 and 3,600 predicted interfaces, respectively, representing 158 and 144 distinct protein–DNA interfaces from the FoldBench benchmark (some of the Boltz-2 calculations failed to complete). The PDB entries are listed in the **Appendix**. Structural accuracy was assessed using DockQv2, calculated with FoldBench code, with thresholds of 0.23, 0.49, and 0.80 used conventionally to distinguish acceptable, medium, and high accuracy predictions. Balanced accuracy (BACC) was calculated with equal weighting of interfaces. We investigated whether the same formulation we used for ligands (constant *d*_0_, aligning on protein, and scoring ligand or nucleic acid, and potentially different cutoffs for AF3 and Boltz-2) would hold for nucleic acids.

In Figure 9, we show kernel density estimates for the *ipSAE*(P→N) score for protein-DNA complexes for AF3 (top) and Boltz-2 (bottom) and values of *d*_0_ = 1,2,3,4,5, and from the standard TM formula with *L* equal to the protein length + nucleic acid length. As with ligands, *ipSAE*(P→N) cutoff values of ∼0.5 and ∼0.8 for accurate protein→DNA structure predictions seem reasonable from the distribution of scores for models with DockQv2 ≥ 0.23 and *d*_0_=2 Å.

**Figure 9.**
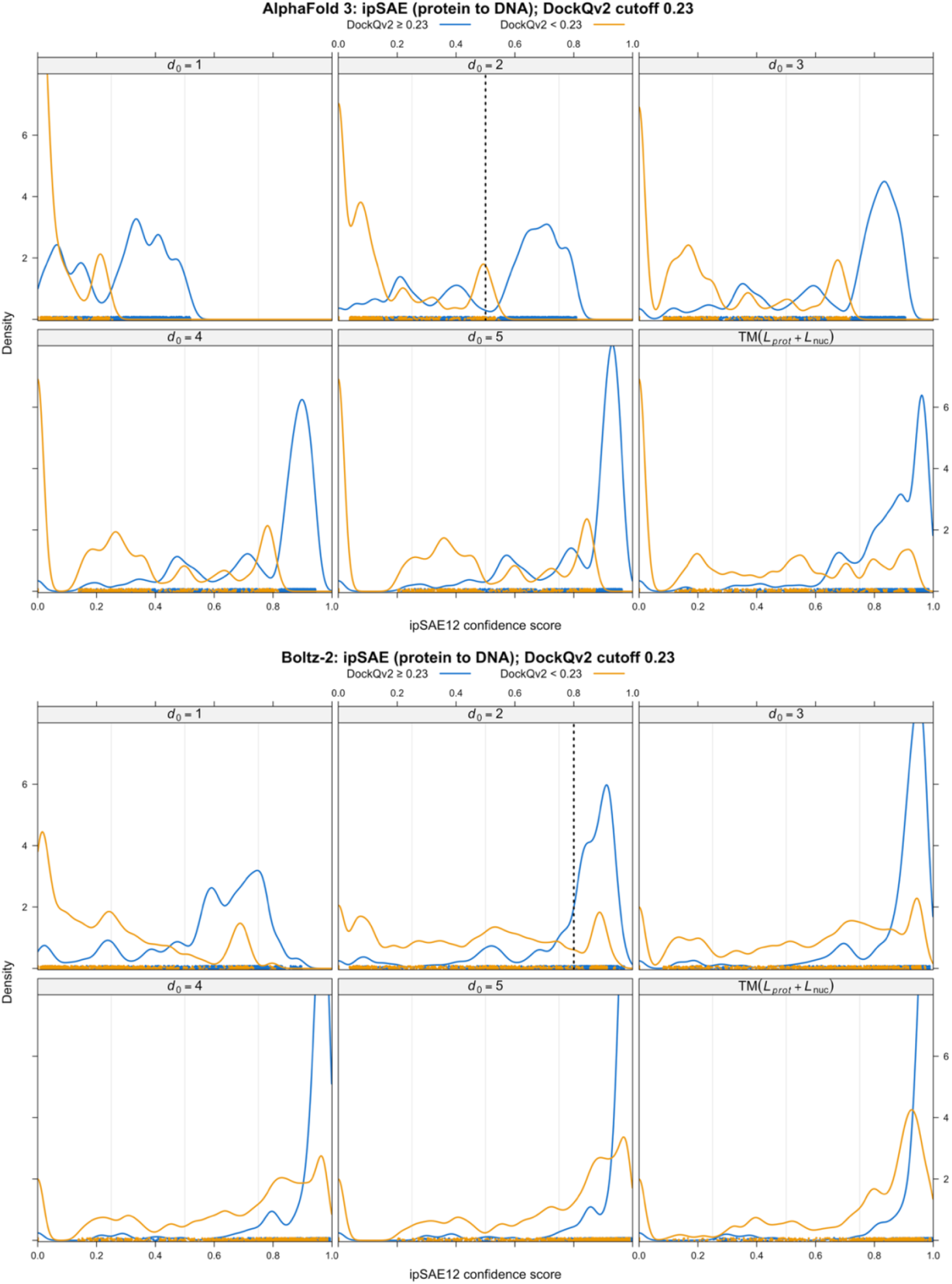
Kernel density estimates of the *ipSAE* score for protein-DNA complexes from FoldBench for AlphaFold3 (top) and Boltz-2 (bottom). Curves are shown for models with DockQv2≥0.23 (blue) and DockQv2<0.23 (orange). The six panels in each plot show the results for d0=1,2,3,4,5 and the TM equation result (as in *ipTM*) with *L* = *L*_*prot*_ + *L*_*nuc*_. Cutoffs for *ipSAE* scores with *d*_0_=2 are shown as dotted lines for AF3 (at 0.5) and for Boltz-2 (at 0.8).

**Figure 10.**
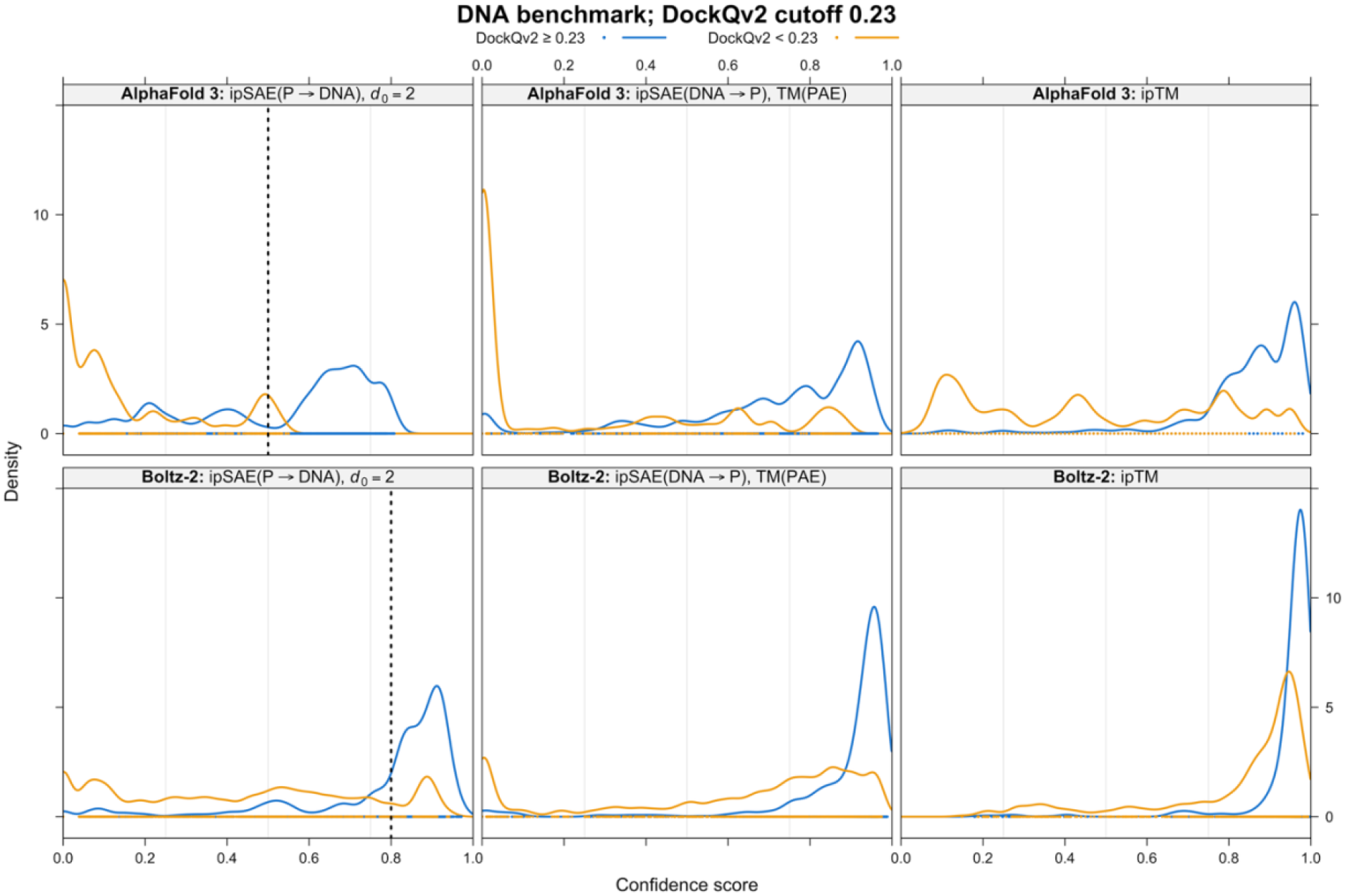
Kernel density estimates of directional *ipSAE* score for protein-DNA complexes from FoldBench for AlphaFold3 (top) and Boltz-2 (bottom). Curves are shown for models with DockQv2≥0.23 (blue) and DockQv2<0.23 (orange). The three panels in each row show the results for *ipSAE*(P**→**N), *ipSAE*(N**→**P), and *ipTM*. Cutoffs for *ipSAE* scores with *d*_0_=2 are shown as dotted lines for AF3 (at 0.5) and for Boltz-2 (at 0.8).

For AF3, protein→DNA *ipSAE* calculated with a fixed *d*_0_ of 2 Å achieved maximum BACC values of 0.812, 0.782, and 0.839 at the three DockQv2 thresholds of 0.23, 0.49, and 0.8, respectively, compared with 0.814, 0.786, and 0.802 for native *ipTM*. At an *ipSAE*(P→N) threshold of 0.5, BACC values were 0.791, 0.767, and 0.825, approaching the maximum values across all three definitions of structural accuracy. For Boltz-2, maximum BACC values for *ipSAE*(P→N) with *d*_0_=2 Å were 0.798, 0.765, and 0.766, compared with 0.789, 0.797, and 0.777 for native *ipTM*. An *ipSAE* threshold of 0.8 yielded BACC values of 0.782, 0.746, and 0.748. In contrast, the optimal native *ipTM* threshold was 0.959 for all three DockQv2 criteria, reflecting the pronounced concentration of Boltz-2 *ipTM* scores near 1.0. It would be likely that a cutoff for *ipTM* for protein-DNA complexes for Boltz-2 would be very sensitive to the benchmark set, given the narrow range and overlap of scores for accurate and inaccurate structures.

Varying fixed *d*_0_ from 1 to 5 Å had little effect on the maximum BACC, although increasing *d*_0_ shifted score distributions toward 1.0. Thus, the choice of *d*_0_ primarily affected the scale and dynamic range of confidence scores rather than their ability to discriminate approximately correct and incorrect predictions. These results support using a fixed *d*_0_ of 2 Å for protein–DNA *ipSAE* as a practical choice that preserves useful score variation while achieving discrimination comparable to native *ipTM*.

#### Protein–RNA complexes

We also evaluated protein–RNA complexes, comprising 27 distinct interfaces predicted by AF3 and 19 by Boltz-2. The RNA dataset was considerably smaller than the DNA dataset and exhibited uneven distributions of DockQv2 scores, limiting comparisons across structural-accuracy thresholds.

For AF3, protein→RNA *ipSAE*(P→N) calculated with *d*_0_ =2 Å achieved maximum BACC values of 0.884 and 0.721 at DockQv2 thresholds of 0.23 and 0.49, respectively, compared with 0.854 and 0.659 for native *ipTM*. For Boltz-2, at a DockQv2 threshold of 0.49, *ipSAE*(P→N) and native *ipTM* achieved similar maximum BACC values of approximately 0.720. However, optimal score thresholds varied between programs and definitions of structural accuracy.

Interpretation of these results was complicated by the limited number of independent RNA complexes and the scarcity of highly inaccurate or highly accurate predictions in some subsets.

In addition, individual complexes showed substantial discrepancies between structural accuracy and predicted confidence. Consequently, these results provide an initial evaluation of directional *ipSAE* for protein–RNA complexes but do not establish an optimal *d*_0_ or a generally applicable confidence threshold. A larger and more diverse benchmark will be needed to determine whether the parameter choices supported by the protein–DNA results also apply to protein–RNA interactions.

## DISCUSSION

In 2025, I introduced the *ipSAE* score to address limitations of the *ipTM* score for protein–protein complexes. Many proteins contain disordered regions or folded domains that do not participate in the interaction between two proteins. Because *ipTM* scores the entirety of one protein relative to alignment frames in the other, these regions can substantially lower the score even when the interacting domains are confidently predicted. *ipSAE* addresses this problem by scoring only residues with *PAE* values below a generous cutoff (typically 10 or 15 Å) and calculating the TM-score scaling parameter, *d*_0_, from the number of residues actually scored, rather than the combined lengths of the proteins. An important practical advantage is that *ipSAE* can be calculated directly from the *PAE* matrix, which is provided by major structure prediction programs, including AF2, AF3, Boltz-2, OpenFold3 (16), Chai-1 (17), and Protenix (18, 19). *ipSAE* has subsequently been used by several groups to evaluate predicted protein–protein interactions (20-22), including the AlphaFold Database team (23).

The limitations of *ipTM* become particularly important when scoring interactions between different types of molecules. For a protein–ligand complex, *ipTM* evaluates the ligand relative to alignment frames in the protein and the protein relative to alignment frames defined by individual ligand atoms, retaining the maximum score. Consequently, the final score may arise from a single ligand-centered alignment frame even when other parts of the ligand are poorly positioned relative to the protein. Yet the question of interest is generally how confidently the entire ligand is positioned in the protein binding site, rather than how confidently some portion of the protein is positioned relative to one atom of the ligand. A second limitation arises from the *d*_0_ parameter, which increases with the number of tokens in the molecular system. AF3 bases this parameter on the combined sizes of the molecular components being scored, while Boltz-2 uses the size of the entire system. Thus, increasing the length of a protein can raise its *ipTM* score even when the predicted relative placement of the ligand and the *PAE* values remain unchanged. This dependence on molecular size compresses the range of *ipTM* scores and complicates the establishment of consistent thresholds for identifying accurate ligand placements.

Extending *ipSAE* beyond protein–protein interactions therefore requires reconsidering three choices: which molecule provides the alignment frames, which tokens contribute to the score, and how *d*_0_ is determined. These choices should reflect the structural question being asked. For protein–protein complexes, the objective is generally to identify regions of one protein that are confidently positioned relative to another, justifying the exclusion of regions with large *PAE* values. For protein–ligand complexes, by contrast, the objective is to evaluate the placement of the entire ligand, making it appropriate to retain all ligand atoms in the score. I therefore propose aligning on protein residues, scoring all ligand heavy atoms without a *PAE* cutoff, and using a fixed *d*_0_ of 2 Å. More generally, for pairs of different molecular types, the proposed *ipSAE* framework aligns on the more complex component and scores the simpler one. For pairs of molecules of the same general type, where no corresponding directional preference can be defined, the maximum of the two asymmetric scores is retained as a symmetric scoring convention. This does not imply that the two molecules are chemically or structurally equivalent.

The protein–ligand benchmarks support these choices. Using 102 complexes from FoldBench, divided into a development set of 20 complexes and a held-out test set of 82 complexes, I found that protein-to-ligand *ipSAE* with *d*_0_ = 2 Å produces a substantially wider dynamic range than native *ipTM* for both AF3 and Boltz-2. Increasing *d*_0_ shifts the scores toward 1.0, reducing their dynamic range. Reference thresholds of 0.5 for AF3 and 0.8 for Boltz-2, selected using the development set, achieved balanced accuracies of 0.938 and 0.862, respectively, on the independent test set for distinguishing ligand placements with RMSD ≤ 5 Å from those with RMSD > 5 Å. By contrast, we found that *ipTM* for AF3 and Boltz-2 required cutoffs of 0.91 and 0.97 respectively. This agrees with the benchmark of Škrinjar et al who established cutoffs of 0.91 and 0.95 for AF3 and Boltz-1 on the Runs N’ Poses data set respectively (24). These results demonstrate that the proposed formulation provides useful, interpretable score ranges and more practical thresholds for evaluating protein–ligand predictions.

Two additional examples illustrate the consequences of these scoring choices. In predictions of Aurora A kinase with cognate and noncognate ligands, aligning on ligand atoms and scoring the protein can produce relatively high confidence scores when only part of a ligand is plausibly accommodated in the binding site. Scoring the entire ligand relative to protein alignment frames better reflects the predicted placement of the complete molecule. In predictions of the PLCD1 PH domain with its ligand, increasing the length of the protein construct resulted in substantially greater protein–ligand *PAE* values and lower *ipSAE* scores, whereas native *ipTM* changed much less. The ligand RMSDs also increased, although the predicted ligand placements remained relatively accurate. Thus, *ipSAE* was more sensitive to the increased predicted uncertainty, but the PLCD1 results also demonstrate that low confidence scores do not necessarily imply inaccurate ligand coordinates. Nevertheless, both directional *ipSAE* scores are provided, since the reverse direction may prove informative in other applications.

The same principles can be applied to protein–nucleic-acid complexes, although the treatment of scored tokens differs from that for ligands. Nucleic acids may contain regions whose positions relative to a binding protein are poorly determined, particularly in large DNA or RNA molecules. As with proteins, a generous *PAE* cutoff can therefore be used to exclude these regions. Benchmarks of protein–DNA complexes support aligning on protein residues, scoring nucleotides that pass the *PAE* cutoff, and setting *d*_0_ = 2 Å. The resulting scores have a useful dynamic range and achieve balanced accuracies comparable to native *ipTM*. The same practical score thresholds of approximately 0.5 for AF3 and 0.8 for Boltz-2 also perform reasonably well. The considerably smaller protein–RNA benchmark produced less consistent results, however, and additional data will be required to determine whether the same parameter choices and thresholds are optimal for RNA.

Several limitations should be considered. First, *ipSAE* measures predicted structural confidence, not binding affinity or experimental evidence that two molecules interact. Second, although a fixed *d*_0_ of 2 Å produces useful score ranges for the ligand and DNA benchmarks examined here, the optimal parameterization and score thresholds may depend on the molecular system and prediction program. Third, *ipSAE* is calculated from the *PAE* matrix, whereas native *ipTM* is calculated from the underlying predicted aligned-error probability distributions, so differences between the two metrics cannot necessarily be attributed solely to alignment direction, token selection, or *d*_0_. Fourth, predicted confidence does not always correspond directly to structural accuracy, as illustrated by the PLCD1 example. Finally, although the framework provides scores for interactions involving atomic ions, ligand pairs, and nucleic-acid–ligand complexes, these applications remain to be systematically validated.

The importance of alignment direction may extend beyond interactions between different molecular types. Antibody-antigen interactions are asymmetric in the sense that the evolutionary constraints on the antigen and antibody are quite different from each other. Recently, Tupikina et al. used the asymmetric *ipSAE* score (aligning on antibody, scoring antigen) to show that *ipSAE* outperformed *ipTM* in distinguishing experimentally supported antibody binders from putative non-binders across six structure-prediction programs tested (25).

In computational protein binder design, the target and designed binder have distinct roles even though both are proteins. The target has an MSA, potential templates, and likely presence in the training data, while the binder is a single sequence and potentially a new fold that is not present in the PDB. *ipSAE* has been extensively used to evaluate predicted target–binder complexes (10, 11, 26), with some studies favoring the minimum of the two asymmetric scores and others favoring the maximum. However, neither convention directly addresses the structural question of interest: how confidently is the designed binder positioned relative to the target? Aligning on the target and scoring the binder may therefore be a more interpretable choice. One possible explanation for the reported utility of *ipSAE_min* is that it may frequently correspond to this direction when the target is larger than the designed binder, although this relationship remains to be tested. Existing datasets of experimentally characterized protein binders could be used to compare the two directional scores with experimental outcomes and determine whether target-to-binder scoring offers an advantage.

More broadly, the results presented here suggest that a single symmetric confidence score is not necessarily the most informative measure of structural accuracy for every molecular interaction. By explicitly choosing the alignment reference, the scored tokens, and the *PAE* scaling parameter, *ipSAE* provides a flexible framework for evaluating the structural relationships of interest in diverse biomolecular complexes.

## CODE and AVAILABILITY

Code for the updated definitions of *ipSAE* is available from https://github.com/DunbrackLab/IPSAE2. The code calculates *ipSAE* values for all molecular pair types and reproduces the values for protein-protein interactions in the first version of *ipSAE*. For other molecule pair types, the definitions of *ipSAE* are presented in this preprint.

The code takes files for the model coordinates, the *PAE* matrix, and optionally summary confidence scores such as *ipTM* from AlphaFold3 and Boltz-2.

~~~
ipsae fold_kras_raf1_gtpmg_model_0.cif fold_kras_raf1_gtpmg_full_data_0.json \ fold_kras_raf1_gtpmg_summary_confidences_0.json
~~~

Alternatively, it can take a directory name, and it will process all the models in that folder (and in subdirectories). This is probably the easiest way to use it. After installation, the program name is `ipsae` and it can be run on a folder in this way:

~~~
ipsae fold_kras_raf1_gtpmg --output-file kras_raft_gtpmg.tsv
~~~

The tab-delimited output looks like this for an example of KRAS, the RAF1 RAS-binding domain, GTP, and a magnesium ion from the AlphaFold3 server (opened in Excel and formatted):

| model | chn1 | chn2 | type1 | type2 | mol1 | mol2 | iptm | ipsae | ipsae12 | ipsae21 | ipsae_source | align_token12 | align_token21 | iPAE12 | iPAE21 | n0_12 | n0_21 | d0_12 | d0_21 | n_scored_contact12 | n_scored_contact21 | pae_cutoff12 | pae_cutoff21 | native_iptm12 | native_iptm21 |
|---|---|---|---|---|---|---|---|---|---|---|---|---|---|---|---|---|---|---|---|---|---|---|---|---|---|
| fold_kras_raf1_gtpmg_model_0.cif | A | B | prot | prot | AA_188 | AA_80 | 0.86 | 0.7663 | 0.6740 | 0.7663 | ipsae21 | ILE:55:CA | THR:17:CA | 2.28 | 2.77 | 79 | 177 | 3.16 | 4.96 | 16 | 16 | 10 | 10 | NA | NA |
| fold_kras_raf1_gtpmg_model_1.cif | A | B | prot | prot | AA_188 | AA_80 | 0.86 | 0.7648 | 0.6694 | 0.7648 | ipsae21 | ILE:55:CA | THR:17:CA | 2.37 | 2.78 | 80 | 177 | 3.19 | 4.96 | 17 | 16 | 10 | 10 | NA | NA |
| fold_kras_raf1_gtpmg_model_2.cif | A | B | prot | prot | AA_188 | AA_80 | 0.86 | 0.7608 | 0.6655 | 0.7608 | ipsae21 | TYR:40:CA | THR:17:CA | 2.34 | 2.82 | 79 | 177 | 3.16 | 4.96 | 17 | 18 | 10 | 10 | NA | NA |
| fold_kras_raf1_gtpmg_model_3.cif | A | B | prot | prot | AA_188 | AA_80 | 0.86 | 0.7578 | 0.6583 | 0.7578 | ipsae21 | ILE:55:CA | PHE:10:CA | 2.44 | 2.84 | 80 | 177 | 3.19 | 4.96 | 17 | 19 | 10 | 10 | NA | NA |
| fold_kras_raf1_gtpmg_model_4.cif | A | B | prot | prot | AA_188 | AA_80 | 0.86 | 0.7555 | 0.6645 | 0.7555 | ipsae21 | ILE:55:CA | PHE:10:CA | 2.33 | 2.85 | 79 | 177 | 3.16 | 4.96 | 16 | 15 | 10 | 10 | NA | NA |
| fold_kras_raf1_gtpmg_model_0.cif | A | C | prot | lig | AA_188 | GTP | 0.97 | 0.8152 | 0.8152 | 0.8710 | ipsae12 | ALA:18:CA | GTP:1:C5 | 0.95 | 1.88 | 32 | 177 | 2.00 | 4.96 | 32 | 31 | none | 10 | NA | NA |
| fold_kras_raf1_gtpmg_model_1.cif | A | C | prot | lig | AA_188 | GTP | 0.97 | 0.8152 | 0.8152 | 0.8715 | ipsae12 | ALA:18:CA | GTP:1:C5 | 0.95 | 1.88 | 32 | 177 | 2.00 | 4.96 | 32 | 31 | none | 10 | NA | NA |
| fold_kras_raf1_gtpmg_model_2.cif | A | C | prot | lig | AA_188 | GTP | 0.97 | 0.8152 | 0.8152 | 0.8678 | ipsae12 | ALA:18:CA | GTP:1:C5 | 0.95 | 1.91 | 32 | 177 | 2.00 | 4.96 | 32 | 31 | none | 10 | NA | NA |
| fold_kras_raf1_gtpmg_model_3.cif | A | C | prot | lig | AA_188 | GTP | 0.97 | 0.8134 | 0.8134 | 0.8659 | ipsae12 | ALA:18:CA | GTP:1:C5 | 0.96 | 1.93 | 32 | 177 | 2.00 | 4.96 | 32 | 31 | none | 10 | NA | NA |
| fold_kras_raf1_gtpmg_model_4.cif | A | C | prot | lig | AA_188 | GTP | 0.97 | 0.8152 | 0.8152 | 0.8677 | ipsae12 | ALA:18:CA | GTP:1:C5 | 0.95 | 1.91 | 32 | 177 | 2.00 | 4.96 | 32 | 31 | none | 10 | NA | NA |
| fold_kras_raf1_gtpmg_model_0.cif | A | D | prot | ion | AA_188 | MG | 0.97 | 0.8316 | 0.8316 | NA | ipsae12 | LYS:16:CA | NA | 0.90 | NA | 1 | NA | 2.00 | NA | 1 | NA | none | NA | NA | NA |
| fold_kras_raf1_gtpmg_model_1.cif | A | D | prot | ion | AA_188 | MG | 0.97 | 0.8316 | 0.8316 | NA | ipsae12 | LYS:16:CA | NA | 0.90 | NA | 1 | NA | 2.00 | NA | 1 | NA | none | NA | NA | NA |
| fold_kras_raf1_gtpmg_model_2.cif | A | D | prot | ion | AA_188 | MG | 0.97 | 0.8316 | 0.8316 | NA | ipsae12 | LYS:16:CA | NA | 0.90 | NA | 1 | NA | 2.00 | NA | 1 | NA | none | NA | NA | NA |
| fold_kras_raf1_gtpmg_model_3.cif | A | D | prot | ion | AA_188 | MG | 0.97 | 0.8316 | 0.8316 | NA | ipsae12 | LYS:16:CA | NA | 0.90 | NA | 1 | NA | 2.00 | NA | 1 | NA | none | NA | NA | NA |
| fold_kras_raf1_gtpmg_model_4.cif | A | D | prot | ion | AA_188 | MG | 0.97 | 0.8316 | 0.8316 | NA | ipsae12 | LYS:16:CA | NA | 0.90 | NA | 1 | NA | 2.00 | NA | 1 | NA | none | NA | NA | NA |
| fold_kras_raf1_gtpmg_model_0.cif | C | D | lig | ion | GTP | MG | 0.67 | 0.7353 | 0.7353 | NA | ipsae12 | GTP:1:C1' | NA | 1.20 | NA | 1 | NA | 2.00 | NA | 1 | NA | none | NA | NA | NA |
| fold_kras_raf1_gtpmg_model_1.cif | C | D | lig | ion | GTP | MG | 0.67 | 0.7353 | 0.7353 | NA | ipsae12 | GTP:1:N9 | NA | 1.20 | NA | 1 | NA | 2.00 | NA | 1 | NA | none | NA | NA | NA |
| fold_kras_raf1_gtpmg_model_2.cif | C | D | lig | ion | GTP | MG | 0.67 | 0.7353 | 0.7353 | NA | ipsae12 | GTP:1:N9 | NA | 1.20 | NA | 1 | NA | 2.00 | NA | 1 | NA | none | NA | NA | NA |
| fold_kras_raf1_gtpmg_model_3.cif | C | D | lig | ion | GTP | MG | 0.67 | 0.7353 | 0.7353 | NA | ipsae12 | GTP:1:N9 | NA | 1.20 | NA | 1 | NA | 2.00 | NA | 1 | NA | none | NA | NA | NA |
| fold_kras_raf1_gtpmg_model_4.cif | C | D | lig | ion | GTP | MG | 0.66 | 0.7353 | 0.7353 | NA | ipsae12 | GTP:1:N9 | NA | 1.20 | NA | 1 | NA | 2.00 | NA | 1 | NA | none | NA | NA | NA |

where the columns refer to molecule 1 (`chn1, type1, mol1`) and molecule 2 (`chn2, type2, mol2`) and the directional scores (e.g., `ipsae12` is the score for aligning on tokens in molecule 1 and scoring tokens in molecule 2). In the mol columns, proteins are given by “AA” plus the length (e.g. `AA_188` for KRAS in this example). The `iptm` is obtained from the summary confidences file from the AlphaFold server. The true ipsae score is given in the `ipsae` column. The source of this score is given in the `ipsae_source` column, which indicates whether the score came from the `ipsae12` or `ipsae21` directional scores. Protein-protein scores come from the max of these two scores (in this example, from `ipsae21`), while protein/ligand and protein/nucleic-acid scores are by definition from the `ipsae12` score. The tokens which produce the `ipsae12` and `ipsae21` scores are given in the align_token12 and align_token21 columns respectively. The `iPAE12` and `iPAE21` give the average *PAE* of the `n0_12` and `n0_21` scored tokens given the alignment tokens listed in the `align_token12` and `align_token21` columns respectively. The last two columns give the asymmetric *ipTM* scores from the chain-pairwise *ipTM* matrix (`native_iptm12, native_iptm21`), if these are provided in the confidence metric files from the co-folding program. Boltz-2 provides this asymmetric matrix, while AF3 does not. Hence, in this example, the asymmetric *ipTM* scores are given as NA.

The full list of options can be given with `ipsae -h`:

~~~
usage: ipsae [-h] [--version] [--pae-key PAE_KEY] [--pae-cutoff PAE_CUTOFF] [--contact-cutoff
CONTACT_CUTOFF] [--fixed-d0 FIXED_D0] [--protein-min-d0 PROTEIN_MIN_D0]
             [--all-pairs] [--output-file OUTPUT] [--no-recursive] [--dry-run] [--write-token-df]
[--write-atom-df] [--write-pae-df] [--write-by-token] [--write-pae-png]
             [--write-interface] [--write-pymol]
             files [files …]
Directional ipSAE2 from coordinates, PAE, and optional native confidence output positional arguments:
files Two or three predictor files in any order, OR a directory containing models
(recursive by default)
options:
  -h, --help show this help message and exit
  --version show program’s version number and exit
  --pae-key PAE_KEY
  --pae-cutoff PAE_CUTOFF
  --contact-cutoff CONTACT_CUTOFF
  --fixed-d0 FIXED_D0 d0 for scored nucleic acids, ligands, and ions (default 2 Å)
  --protein-min-d0 PROTEIN_MIN_D0
                         Minimum d0 for scored proteins (default 1 Å)
  --all-pairs            Include all molecular pairs, even if not in contact
  --output-file, -output_file, --output OUTPUT
                         Single-model TSV path, or combined TSV path for directory mode
  --no-recursive         When given a directory, scan only that directory
  --dry-run              List model/PAE/confidence matches in directory mode without scoring
  --write-token-df
  --write-atom-df
  --write-pae-df         Large complete interchain directional token-pair PAE table
  --write-by-token       Detailed scores for every eligible alignment token
  --write-pae-png
  --write-interface      Write contacting-residue selections
  --write-pymol          Write PyMOL selections for contacting residues
~~~

The `--pae-cutoff PAE_KEY` argument provides the key name in the json or npz file for the *PAE* matrix. This is not needed for AF3 or Boltz-2 output but might be needed for other programs. The contact cutoff only affects the number of residues listed under `n_scored_contacts12` and `n_scored_contacts21`; it does NOT affect the *ipSAE* scores. Several data frames can be output with options `–write-token-df, --write-atom-df, --write-pae-df, --write-by-token`, and `--write-interface`. A graphical image of the PAE matrix can be output for each model with `–write-pae-png` and a PyMOL script for coloring chains by contacting residues with good PAE scores can be output with `–write-pymol`.

## ACKNOWLEDGMENTS

This work was funded by NIH Grants R35 GM122517 (R. L. D.) and P30 CA006927 (Fox Chase Cancer Center). I would like to thank my lab members, Drs. Qifang Xu, Pragya Priyadarshini, and Joan Gizzio for many useful conversations and for testing the code.

## Appendix. PDB entries in the benchmark datasets

PDB entries are listed in alphanumeric order. All targets correspond to biological assembly 1 in the FoldBench dataset; the suffix “-assembly1” is omitted below.

### Dev20_set (20 PDB entries)

7FUM, 7FUO, 7UYB, 8BIQ, 8CEX, 8HKB, 8OV7, 8PA5, 8PGC, 8PGZ, 8PH1, 8PYI, 8QHG, 8QK9, 8SHJ, 8U37, 8U8J, 8USS, 8WCH, 8WKG.

### Test82_set (82 PDB entries)

7EKA, 7FTI, 7FTP, 7FU2, 7FU4, 7FUG, 7FUL, 7FUQ, 7UAD, 7UDQ, 7UYA, 7UYC, 7X6T, 7YL1, 7ZW8, 8ATL, 8ATN, 8AZC, 8BM2, 8BMV, 8BPV, 8BR6, 8BSA, 8C0Q, 8C3U, 8CC4, 8CEY, 8CHI, 8CJK, 8DI4, 8DJE, 8DMT, 8DVW, 8ERD, 8F8E, 8FF8, 8GKF, 8H7M, 8HY1, 8ID0, 8JJT, 8JNY, 8OHU, 8OKE, 8OKG, 8OKJ, 8OKO, 8OKP, 8OLF, 8OUV, 8OYQ, 8P04, 8P08, 8P3D, 8P5I, 8P92, 8P95, 8P9P, 8P9T, 8PB2, 8PG6, 8PG9, 8PGG, 8PGK, 8PGT, 8PGX, 8PH3, 8PQG, 8PYK, 8PYM, 8QTG, 8QTJ, 8QTK, 8RCW, 8RHE, 8SBT, 8SX2, 8U8K, 8UV0, 8UXS, 8WLB, 8YIK.

### Protein–DNA prediction set (96 PDB entries)

7FTN, 8C41, 8C56, 8C58, 8C59, 8FAY, 8G8H, 8G8J, 8HCL, 8IJP, 8IK5, 8ILD, 8ILF, 8ILG, 8ILH, 8IPP, 8IYQ, 8J71, 8JKK, 8K86, 8K8A, 8K8C, 8K8D, 8KFR, 8OK9, 8P5Q, 8PI8, 8PM5, 8PM7, 8PMC, 8PMF, 8PMV, 8PNA, 8PNC, 8PW0, 8Q5O, 8Q9N, 8QMB, 8R0S, 8RM6, 8RM7, 8RPX, 8RZV, 8SLN, 8TAC, 8TLF, 8TLG, 8TLH, 8U0O, 8U0P, 8U6X, 8U8L, 8UFK, 8UJX, 8URK, 8UVK, 8UZT, 8V7B, 8V7G, 8VDH, 8VDN, 8VF8, 8VF9, 8VFC, 8VFD, 8VFE, 8VFF, 8VFG, 8VFJ, 8VMV, 8VZM, 8WQ5, 8WT6, 8X5V, 8XA9, 8XAS, 8YEY, 8YFI, 8YFJ, 8YRS, 8YXW, 8YZS, 8Z2N, 9BDU, 9BDV, 9BDW, 9BS2, 9BVD, 9C3S, 9C3T, 9CI9, 9CIH, 9DP2, 9ENQ, 9EZZ, 9F0E.

AlphaFold3 predictions were generated for all 96 protein–DNA entries. Boltz-2 predictions were generated for 90 of them; the six without Boltz-2 predictions were: 8C41, 8IYQ, 8QMB, 8WT6, 8YRS, 9ENQ. The input sets contained 164 protein–DNA interfaces for AlphaFold3 and 148 for Boltz-2, corresponding to 4,100 and 3,700 interface–model combinations, respectively. After matching to ipSAE scores and DockQv2 evaluations, 158 AlphaFold3 interfaces (3,950 models) and 144 Boltz-2 interfaces (3,600 models) were retained. Thus, the 96-entry DNA list describes the prediction targets, including entries for which matched analyses were unavailable.

## Notes

https://github.com/DunbrackLab/IPSAE2

### Competing Interest Statement

The authors have declared no competing interest.

